# Leveraging targeted sequencing for non-model species: a step-by-step guide to obtain a reduced SNP set and a pipeline to automate data processing in the Antarctic Midge, *Belgica antarctica*

**DOI:** 10.1101/772384

**Authors:** Vitor A. C. Pavinato, Saranga Wijeratne, Drew Spacht, David L. Denlinger, Tea Meulia, Andrew P. Michel

**Author notes:** Corresponding author: Vitor A. C. Pavinato, Department of Entomology, Thorne Hall, CFAES Wooster Campus, The Ohio State University, 1680, Madison Avenue, Wooster, OH, USA.

## Abstract

The sequencing of whole or partial (e.g. reduced representation) genomes are commonly employed in molecular ecology and conservation genetics studies. However, due to sequencing costs, a trade-off between the number of samples and genome coverage can hinder research for non-model organisms. Furthermore, the processing of raw sequences requires familiarity with coding and bioinformatic tools that are not always available. Here, we present a guide for isolating a set of short, SNP-containing genomic regions for use with targeted amplicon sequencing protocols. We also present a python pipeline--PypeAmplicon-- that facilitates processing of reads to individual genotypes. We demonstrate the applicability of our method by generating an informative set of amplicons for genotyping of the Antarctic midge, *Belgica antarctica*, an endemic dipteran species of the Antarctic Peninsula. Our pipeline analyzed raw sequences produced by a combination of high-multiplexed PCR and next-generation sequencing. A total of 38 out of 47 (81%) amplicons designed by our panel were recovered, allowing successful genotyping of 42 out of 55 (76%) targeted SNPs. The sequencing of ∼150 bp around the targeted SNPs also uncovered 80 new SNPs, which complemented our analyses. By comparing overall patterns of genetic diversity and population structure of amplicon data with the low-coverage, whole-genome re-sequencing (lcWGR) data used to isolate the informative amplicons, we were able to demonstrate that amplicon sequencing produces information and results similar to that of lcWGR. Our methods will benefit other research programs where rapid development of population genetic data is needed but yet prevented due to high expense and a lack of bioinformatic experience.

## Introduction

Sequencing whole (Prado-Martinez *et al.*, 2013) or partial (*e.g.* RADseq, Baird *et al.*, (2008)) genomes are now standards in molecular ecology and conservation genetic research (Ekblom & Galindo, 2010; Fuentes-Pardo & Ruzzante, 2017; Supple & Shapiro, 2018). Although sequencing costs per sample and per base-pair are decreasing, expenses to generate sufficient genotypic data still impose serious constraints on the number of individuals or populations sampled (Larson *et al.*, 2019). To estimate important features such as genetic diversity, population structure and selection, genotypes from many individuals and populations provide more robust results (Fumagalli, 2013). A sizable number of individuals must be genotyped regardless of the constraints imposed by the sequencing technology and the available budget. Adequate sampling is particularly important for conservation genetic studies that require the correct delimitation of the targeted taxon for protection (Mace, 2004) and for genetic monitoring. In addition, data generated with next-generation sequencing requires massive computational storage and considerable training in bioinformatics and processing to make genotypic data available for analysis. A lack of technological training restricts the choice of molecular marker systems for laboratories researching conservation genetics (Fuentes-Pardo & Ruzzante, 2017; Taylor, Dussex & van Heezik, 2017) and may require bioinformatic processing by another laboratory or private company, adding to the expense.

Most important information for population and conservation genetic studies can be achieved with traditional molecular methods that do not require whole genome sequencing of populations (Allendorf, 2017; Bowden *et al.*, 2012; Fischer *et al.*, 2017; Peterson *et al.*, 2012). Targeted enrichment protocols are alternatives to whole-genome re-sequencing (WGR) (Henriques *et al.*, 2018; Meek & Larson, 2019; Milano *et al.*, 2013). These protocols enable amplification of specific genomic regions that contain previously discovered variation. Combined with next-generation sequencing and robust multiplex PCR amplification, they allow rapid sequencing of hundreds of regions of the genome of several individuals (Campbell *et al.*, 2014; Yang *et al.*, 2016). Unlike exon capture protocols, which only sequence targeted expressed genes, amplicon sequencing allows sequencing of known regions that likely contain more neutral variation needed for population diversity studies. They also allow improved control and uniformity of sequencing coverage and the acquisition of reliable, genotypic information with a limited constraint on the number of sequenced individuals. These features make amplicon sequencing a valuable tool for population and conservation genetics that require the genotyping of several individuals from multiple populations to accurately estimate neutral genetic diversity, define populations and infer important demography features (*e.g.* isolation-by-distance, effective population size).

As climate change and expansion of extreme environments continue to encroach on ecological communities, researchers will need more rapid and cost-effective methods to assess changes in population and species diversity. Molecular ecology studies of species that already inhabit extreme environments can serve as a model for adaptation and have shown the importance of genetic variation and structure for population persistence (Brown *et al.*, 2019). To completely understand and predict a species’ and/or populations’ potential for extinction, it is necessary to not only uncover the molecular basis of adaptation, but also to characterize past and current responses to environmental change. This is possible by measuring the impact of selection and adaptation relative to the overall genetic diversity and population size to gain an understanding of past and recent demographic changes. In some cases, these studies may require data from fragmented populations, or among species with wide but patchy distributions. Since cost often prohibits whole genome sequencing in populations, targeted sequencing is a less expensive alternative that combines the robustness of genotyping with the practicality of processing large number of samples.

The Antarctic midge, *Belgica antarctica* (Diptera: Chironomidae), is an endemic insect from the Antarctic continent ranging from southern Marguerite Bay (ca. 68°S) northwards to the South Shetland Islands (ca. 63°S) (Convey & Block, 1996). *Belgica antarctica* is an ancient lineage that diverged from the closest Orthocladiinae taxon inhabiting Patagonia, ∼68Myr ago (Allegrucci *et al*., 2006). Populations of *B. antarctica* that we examined near the Palmer Research Station area may have originated from relictual populations that survived continental glaciation or possibly from immigrants arriving from more northerly refuges. The midge exhibits a plethora of adaptations to survive the extreme Antarctic conditions. Loss of wings is a likely adaptation for surviving on the windy off-shore islands (Kelley *et al.* 2014), and numerous physiological adaptations are evident, including freeze tolerance, constitutive expression of heat shock proteins (Rinehart *et al.*, 2006), and resistance to dehydration (Hayward *et al*., 2007; Teets *et al.*, 2012a). By evolving physiological and morphological adaptations to inhabit Antarctica over several million years, this species is highly adapted to surviving under these environmental conditions. However, these conditions are rapidly changing and threaten its persistence.

Here we present a method to obtain an informative and reduced set of SNP markers for targeted enrichment sequencing, and a python pipeline, PypeAmplicon (Wijeratne & Pavinato, 2018), designed to facilitate processing of raw amplicon reads produced by a combination of high-mutiplexed PCR and short read sequencing. Starting with low coverage, whole-genome re-sequencing data (lcWGR), we present the steps to isolate informative and robust markers, with guidelines for marker filtering based on population genetic estimates. This method provided an informative set of amplicons and SNPs for *B. antarctica*. We present some helpful guidance on how to process the raw data produced by a multiplex-PCR based amplicon sequencing for rapid and reliable genotyping. By comparing the summary statistics estimated with the lcWGR data with that obtained with the new amplicons set, we show that the platforms produced similar patterns of genetic diversity and population structure.

## Material and Methods

### Panel design for target sequencing enrichment

#### Biological material and DNA extraction

For the whole-genome re-sequencing we sampled individuals from two sites that were 7.8 km apart: Humble Island (HP) and Dream Island (D1), in the Antarctic Peninsula (Figure S1). Genomic DNA from twelve adult individuals from each site were extracted using the DNeasy® Blood & Tissue Kit (Qiagen). DNA was eluted in 50 µl of TE buffer (10 mM Tris-HCl pH 8.0 and 1 mM de EDTA pH 8.0) and stored at −20°C. Prior to library preparation, DNA samples were quantified using a Qbit® kit (Invitrogen). The instrument was calibrated for the Quant-iT dsDNA BR Assay (assay range between 2–1000 ng; starting sample concentration between 100 pg/µl and 1µg/µl), and samples were prepared according to the manufacturer’s instructions. DNA samples were diluted with ddH_2_O to reach our target concentration of 50ng/uL.

#### De novo sequencing for marker discovery

For each sample, we obtained a whole-genome re-sequencing library using the TruSeq Library Prep Kit (Illumina). For each individual, one unique barcode was added to the 5′ end, allowing us to recover short reads from each sample after parallel sequencing. Samples were pooled and sequenced with the Hiseq® 2500 System (Illumina). Paired-end sequencing with 100 cycles for each side of the fragments was performed in one lane.

#### Marker discovery, genetic diversity, population structure and outlier detection

SNP discovery and genotyping were carried out following a reference-based pipeline (Figure S2). Trimmomatic (Bolger *et al.*, 2014) was used to remove low quality reads and nucleotides, any remaining Illumina barcodes, and adapters. For each round of quality control, the quality of fastq reads was checked with FASTQC (http://www.bioinformatics.babraham.ac.uk/projects/fastqc/). Neither Miseq adapter nor kmer content were observed after trimming. Reads that passed quality control were aligned to a reference genome (Kelley *et al.* (2014); found at NCBI BioProject <u>#PRJNA172148</u>) using Bowtie2 (Langmead & Salzberg, 2012). SAMtools (Li *et al.*, 2009) was used to convert SAM to BAM files and to call SNPs. We applied a stringent filter with vcftools version 0.1.16 (Danecek *et al.*, 2011) to keep only biallelic SNPs with a GQ quality higher than 20, an average depth between 30 and 80 reads, and that were present in at least 80% of the individuals with minor allele frequency higher than 0.001.

Global and within-population genetic variation were summarized by calculating the folded allele-frequency spectrum (AFS) including the observed and expected heterozygosity. The folded AFS was obtained with a custom R script using the matrix of SNPs produced by vcftools version 0.1.16, where 0 is the reference allele, 1 corresponds to the heterozygous genotypes and 2 is the alternative allele. The proportion of heterozygous genotypes (H_O_ and H_E_) were calculated with R package adegenet version 2.1.1 (Jombart, 2008). Fisher’s exact tests for the H-W proportion were carried out for all filtered SNPs in each population with 1000 Monte Carlo permutations using the R package *pegas* version 0.11 (Paradis, 2010). P-values were corrected by Bonferroni’s procedure. The population genetic structure was summarized with global F_ST,_ estimated with the R package hierfstat version 0.04-22 (Goudet, 2005) along with a principal component analysis (PCA) carried out with adegenet.

We checked the neutrality of all whole-genome re-sequencing SNPs (*“WGR SNPs”*) by running two genome scan analyses based on the distribution of F_ST_: BAYESCAN v2.1 (Foll & Gaggiotti, 2008) and OUTFLANK (Whitlock & Lotterhos, 2015). For BAYESCAN, the analysis included 500,000 Markov-Chain Monte-Carlo (MCMC) after 500,000 iterations of a burn-in phase. We considered SNPs as outliers if they had posterior intro-locus F_ST_ estimates higher than the upper limit of the 95% highest posterior density (HPD) interval. For OUTFLANK, we used a set of independent SNPs to calibrate the F_ST_ null distribution and considered outlier SNPs with a q-value lower than 0.10, thus being less conservative with a false discovery rate (FDR) of 10%. A set of independent SNPs were identified by only taking SNPs with intra- and inter-scaffold R^2^ statistics lower than 0.01. R^2^ statistics were calculated with vcftools (Danecek *et al.*, 2011). We only considered a SNP an outlier if it was identified in both analyses. We assessed if an outlier SNP was within or near a gene by visually mapping back to the genome and annotated gene list. We also determined if the variant alleles impacted predicted genes (i.e. nonsynonymous) by running a variant effect prediction analysis in Ensembl metazoa (https://metazoa.ensembl.org/index.html).

#### Isolation of candidate markers and panel design

To reduce the WGR SNPs to an informative and reduced set of SNPs, we performed a PCA with all discovered SNPs and ranked the first principal component loading score (Hulsegge *et al.*, 2013; Wilkinson *et al.*, 2011). PCA loadings were obtained with R package adegenet (Jombart, 2008). Since PCA loadings represent a correlation between the SNP and the respective principal components (PCs), ranking the first PC loadings reveals SNPs that contributed most to the individual assignment. Selected SNPs were only included in the panel if they met all the required characteristics: 1) passed the within population Fisher’s exact test of Hardy-Weinberg equilibrium; 2) were not heterozygous in all sequenced individuals (as the heterozygote excess could be due to duplication or errors in SNP allele calling); and 3) were at least 5bp apart from another SNP in the scaffold, as clusters of SNPs could indicate accumulation of errors during sequencing and alignment of reads. By removing SNPs with only heterozygous states, in disequilibrium or in clusters, we improved the quality of SNP selection and minimized impact of low coverage sequencing on marker selection.

When possible, WGR SNPs detected as outliers were retained since their status can be validated with genotyping of additional populations and individuals. We manually inspected SNPs in annotated genes of the *B. antarctica* genome (Kelley *et al.*, 2014) that may also be adaptive and added those that were polymorphic. We also included randomly chosen neutral SNPs in the panel, since the choice based on ranked PCA can be biased towards SNPs that are divergent (neutrally or linked to a selected locus) between populations. A total of 47 primer pairs were designed to amplify 57 targeted SNP markers that included neutral and outlier SNPs, and SNPs within genes (Figure S3). Each primer pair amplified a region ∼150 bp long and was manufactured by Fluidigm® according to the Access Array^™^ system. We ensured that each primer uniquely paired with its corresponding amplicon sequence and if the amplicon aligned uniquely to itself. We also checked if each amplified region aligned to a unique region in the genome, by performing a homology search with BLAST (Altschul *et al.*, 1990).

### Validation of the panel for target sequencing enrichment

#### Samples

To validate the isolated candidate SNPs, we genotyped, whenever possible, the same individuals used in the whole-genome re-sequencing. Total genomic DNA was already extracted (see above), but only 21 out of 24 adults (11 from D1 and 10 from HP population) had enough DNA remaining to be quantified using NanoDrop® (ThermoFisher). DNA samples were normalized (when necessary) to reach a target concentration of 15 ng/uL for analysis on the Fluidigm Access Array (FAA).

#### Targeted resequencing

Amplicon sequencing was carried out following the FAA system protocol that automates preparation of amplicon-based libraries for up to 48 samples. Primers were designed to amplify specific regions of the genome spanning 150 bp that contained the isolated targeted SNPs. The multiplex PCR libraries were prepared following the Fluidigm® library prep 48.48 IFC protocol, which consisted of two amplification steps: 1) a primary PCR that amplified the amplicon of each targeted region, and 2) a secondary PCR on the pool of amplicons from each sample to attach an individual-specific barcode. All samples were then pooled and the NGS library was prepared following the NEBNext® UltraTM DNA Library Prep Kit for Illumina® (New EnglandBioLabs Inc.). Paired-end sequencing with 150 cycles for each side of the fragments was performed in one lane of the MiSeq® System (Illumina).

#### SNP and genotyping calling for amplicon sequencing

The raw paired-end reads of each sample were first merged and clustered by similarities (> 75%) with usearch (Edgar, 2010). Orphaned reads and clusters with fewer than 25 reads were discarded. Reads that passed the clustering filter were then split into different files with BBsplit, according to their similarity to each reference amplicon. This served a dual purpose: 1) to split the reads by amplicon, and 2) as a quality control for reads within clusters. Filtered reads were then aligned to their amplicon using BBmap. Both BBsplit and BBmap are part of BBtools (https://sourceforge.net/projects/bbmap/). Each multi-sample BAM file of each amplicon was then merged.

We developed a pipeline to automate the processing of Fluidigm/MiSeq raw reads to call individual genotypes. The pipeline PypeAmplicon is available at zenodo (doi:10.5281/zenodo.1490421). The pipeline was developed to process targeted-enrichment amplicon-sequencing produced by double PCR protocols such as the FAA, but it can work with raw data produced by any amplicon-sequencing protocol as long as the reference amplicon sequences are provided (fasta format). The pipeline was designed as an alternative to the alignment of filtered reads to a reference genome, since most non-model species may not have a sequenced genome. The outputs of the pipeline include BAM files so that any SNP caller can be used such as GATK (McKenna *et al.*, 2010), SAMtools (Li *et al.*, 2009), or ANGSD (Korneliussen *et al.*, 2014) (the latter is preferred if the goal is to obtain genotype-likelihoods with limited sequencing output). In this study, we used Freebayes (Garrison & Marth, 2012) to perform the SNP calling on the multi -sample and -amplicons BAM file (Figure S4). To facilitate comparison of the SNPs and genotypes produced by the whole-genome and amplicon sequencing, the parameter that controls size of haplotype gaps (-E) was set to one. We only kept SNP variation (local haplotypes were discarded) and produced two datasets for downstream analyses and comparisons: 1) only targeted SNPs that were recovered and 2) all SNPs, also called *“Amplicon-seq SNPs”*.

#### Validation of targeted SNPs and comparison between WGR and amplicon-seq SNPs

We performed two types of comparisons to validate the targeted SNP approach. First, we compared only the genotypes and the proportion of missing data from the targeted SNPs present in the whole-genome sequencing and in the amplicon-sequencing data (i.e. same data but different technologies). This step allowed us to evaluate correspondence between genotypes called with both sequencing approaches. Assuming the amplicon-sequencing protocol provides better confidence in genotyping calls since it allows sequencing of targeted regions with more depth, we expected to have less missing calls for each individual and targeted SNP. We also expected a reduction in the proportion of homozygous genotypes miscalled as heterozygous (false positives) and in the proportion of heterozygous genotypes miscalled as homozygous (false negatives). This comparison also allowed us to globally evaluate the isolation of informative SNPs and validate genotypes identified by the outlier approaches.

For the second comparison, we evaluated the ability of the amplicon-sequencing approach to recover similar estimates of population genetic summary statistics and degrees of population allele frequency differences through the quantification of overall F_ST_ and genetic structure. For this comparison, we used two datasets: one containing all SNPs discovered with whole-genome sequencing (WGR SNPs) and one containing all SNPs (originally targeted plus the *de novo* SNPs) obtained with amplicon-sequencing. In addition to estimates obtained for the WGR SNPs (see above), we also calculated summary statistics including: number of nucleotide differences per nucleotide site (*π*), Tajima’s D (Tajima, 1989) and Watterson’s estimator (Watterson, 1975). Estimates were performed for each focal SNP and a 150bp window around the SNP (to reproduce average size of the amplicon-sequencing fragment). Estimates were obtained for each population and globally using egglib v3.0.0b21 (Mita & Siol, 2012).

## Results

### Panel design for target sequencing enrichment

#### Marker discovery, genetic diversity, population structure and outlier detection

We re-sequenced the whole-genome of twenty-four individuals of *B. antarctica* collected from two islands of the Antarctic Peninsula. A total of 196,799,812, 100bp-long paired-end reads were produced, corresponding to an average coverage of 16X per individual. Sequencing data of one individual, from Dream Island, was removed from the downstream analysis due to low yield. We applied a stringent quality control on the raw reads and on the called SNPs to minimize accumulating sequenc errors during steps of the reduced SNP panel design. Coverage was reduced to 13X and 11X after trimming and mapping (Figure S5A). We initially found 715,721 variants including SNPs and INDELs. The stringent filter reduced the variants to 1,260 SNPs with an mean coverage of ∼50X (Figure S5B). The final set of filtered SNPs contained only bi-allelic SNPs, with a PHRED quality score > 20 and an average 8.53% of missing data (with a maximum of 17.39%). We limited minimum and maximum depth and the missing data to minimize miscalling of heterozygous genotypes.

With the WGR SNPs, we were able to characterize within population allele frequencies using SNP markers and assess the population genetic structure with individual genotype data. The folded allele-frequency spectrum (AFS) showed an excess of intermediate-frequency variants for each population (Figure 1A). Since the folded AFS ranged from 1 to *n* number of alleles for each site (or 1/2*n* to *n*/2*n*) with *n* being the number of haploid genomes, the rightmost part of the AFS represents the number of loci that have an allele frequency equal to or close to 0.5. The p-values of Fisher’s exact test were calculated globally (pooling the two populations) and for each population to evaluate the deviation of H-W proportions. The Q-Q plot showed the majority of loci deviated from the expected distribution of p-values. Globally, 57.03% of the loci (and ∼55% in each population) deviated from the H-W proportions when no correction for multiple tests was applied (Figure S6). With a Bonferroni’s correction, 37.03% globally deviated from the H-W proportion. The proportion of heterozygous genotypes observed globally and in each population were high: 0.616 globally; 0.715 in D1 and 0.739 in HP, compared to the expected proportion of 0.344 globally; 0.395 in D1 and 0.404 in HP. We calculated within- and between-scaffolds R^2^ and estimated LD decay and the population recombination rate (ρ = 4Nr) using the Weir & Hill equation (Weir and Hill 1988). When populations were combined, the distance of half LD decay was 476bp, and for D1 and HP populations, half decay distances were 1072bp and 410bp (Figure S7). Population recombination rates, ρ, were ∼4e-3 globally, and ∼2e-3 and ∼6e-3 for D1 and HP, respectively. Overall genetic structure measured with global F_ST_ was 0.028 (with bootstrap 95% confidence interval ranging from 0.022 to 0.032), and PCA showed two major clusters on each side of the first PC that represented each population (Figure 1B).

**Figure 1.**
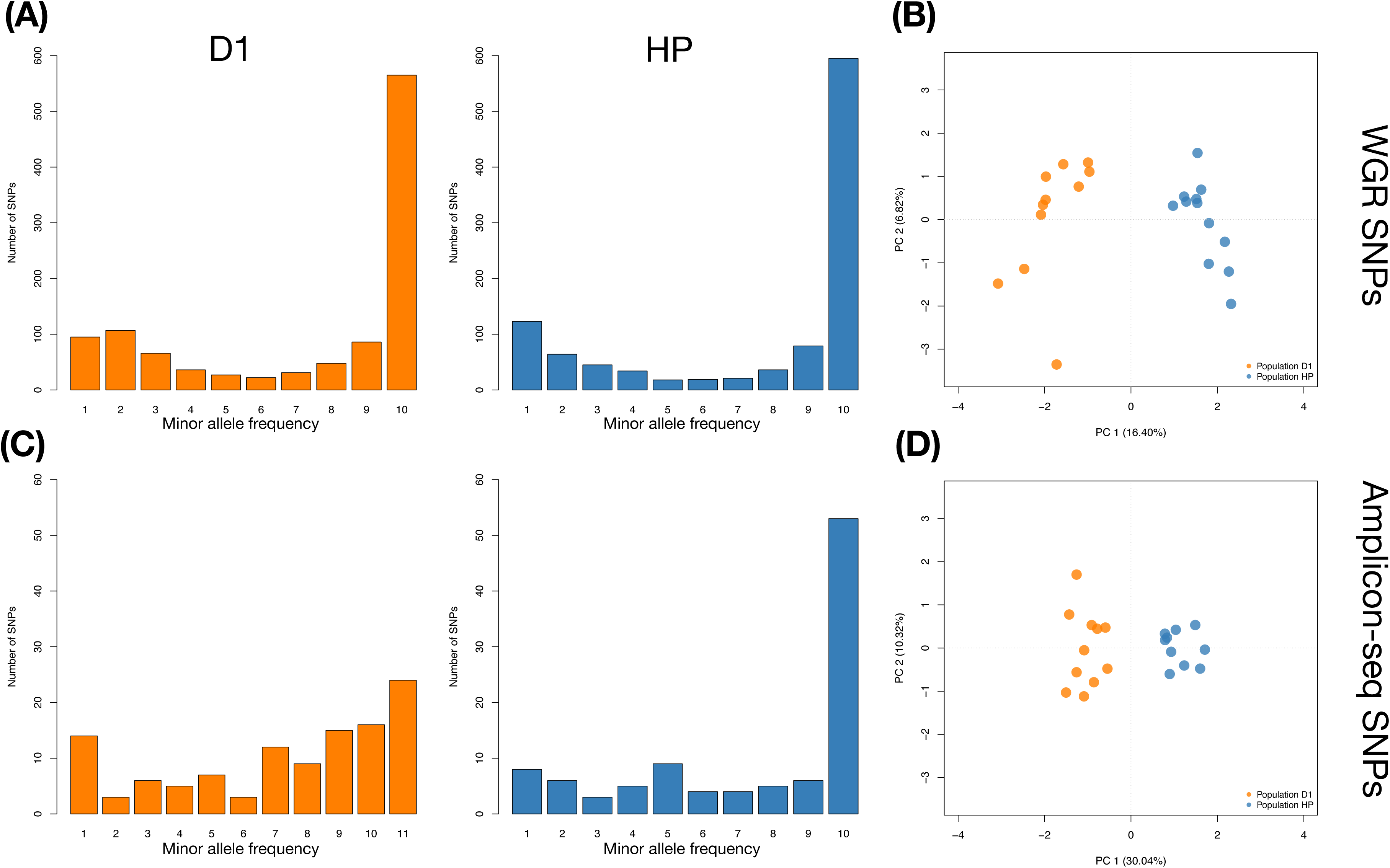
Folded AFS and PCA for lcWGR and amplicon-sequencing SNPs. A) Folded-AFS summarizing the allele frequency distribution of SNPs obtained with lcWGR. B) PCA for lcWGR. C) the folded-AFS of SNPs obtained with amplicon-sequencing. D) PCA for amplicon-sequencing.

The Bayesian method implemented in BAYESCAN did not find significant outliers even with a FDR of 10%. Since an excess of heterozygous loci in our SNP data can indicate a deviation in the island model, and can bias the expected global F_ST_ values, we applied a more flexible criteria to accept a locus as an outlier. We found eight outliers by considering loci that had a posterior F_ST_ value higher than the upper 95% HPD interval. The method implemented in OUTFLANK found 31 outliers. We found 8 outlier loci common to both analyses (Figure 2). The outlier SNP with the highest posterior F_ST_ was found inside a gene that encodes a putative vitellogenin protein where the alternative allele is a non-synonymous mutation. The second outlier SNP found in the same gene had a lower posterior F_ST_, but both SNPs had contrasting values. It is likely this outlier was polymorphic in D1, but was fixed for the reference allele in HP. The second highest posterior F_ST_ belonged to an uncharacterized protein, and the alternative allele was a synonymous mutation (Table S1).

**Figure 2.**
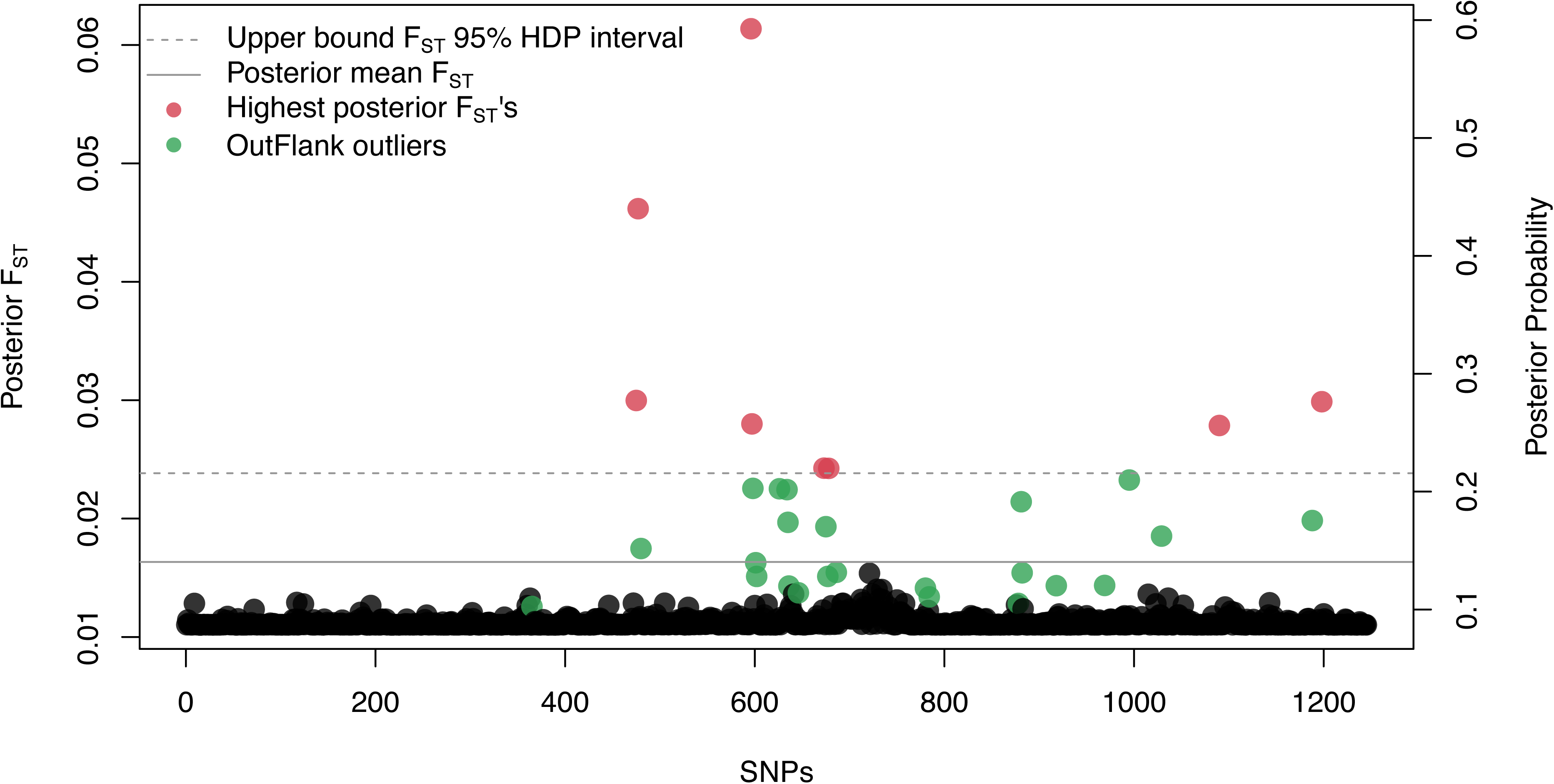
F_ST_-based outlier detection analysis. Green represents SNPs detected as outliers by OUTFLANK and in pink are the SNPs with posterior F_ST_ higher than the upper limit of the 95% F_ST_ highest posterior density (HDP) interval.

#### Isolation of candidate markers and panel design

Third-eight SNPs were identified as informative after ranking PCA scores. Four of these SNPs were previously identified as outliers with the highest posterior F_ST_; twelve were identified inside an open-reading frame, but without functional information. Additionally, ten randomly chosen were added (Figure 3A). We included four SNPs in genes that may be associated with a physiological mechanism to survive in Antarctica: one SNP in the gene *PEPCK* (associated with cold and drought tolerance, Teets *et al.*(2012b)); one SNP in *Buffy* which regulates cell death in *D. melanogaster* (Danial & Korsmeyer, 2004; Quinn, 2003); one SNP in the ecdysone receptor *EcR* (Hill *et al.*, 2013), and a SNP in the RNA-interference machinery *Dicer* (Dicer, Lee *et al.* (2004)). In total, 55 SNPs in 47 amplicons were targeted by the panel. Primers were designed following the FAA system recommendations.

**Figure 3.**
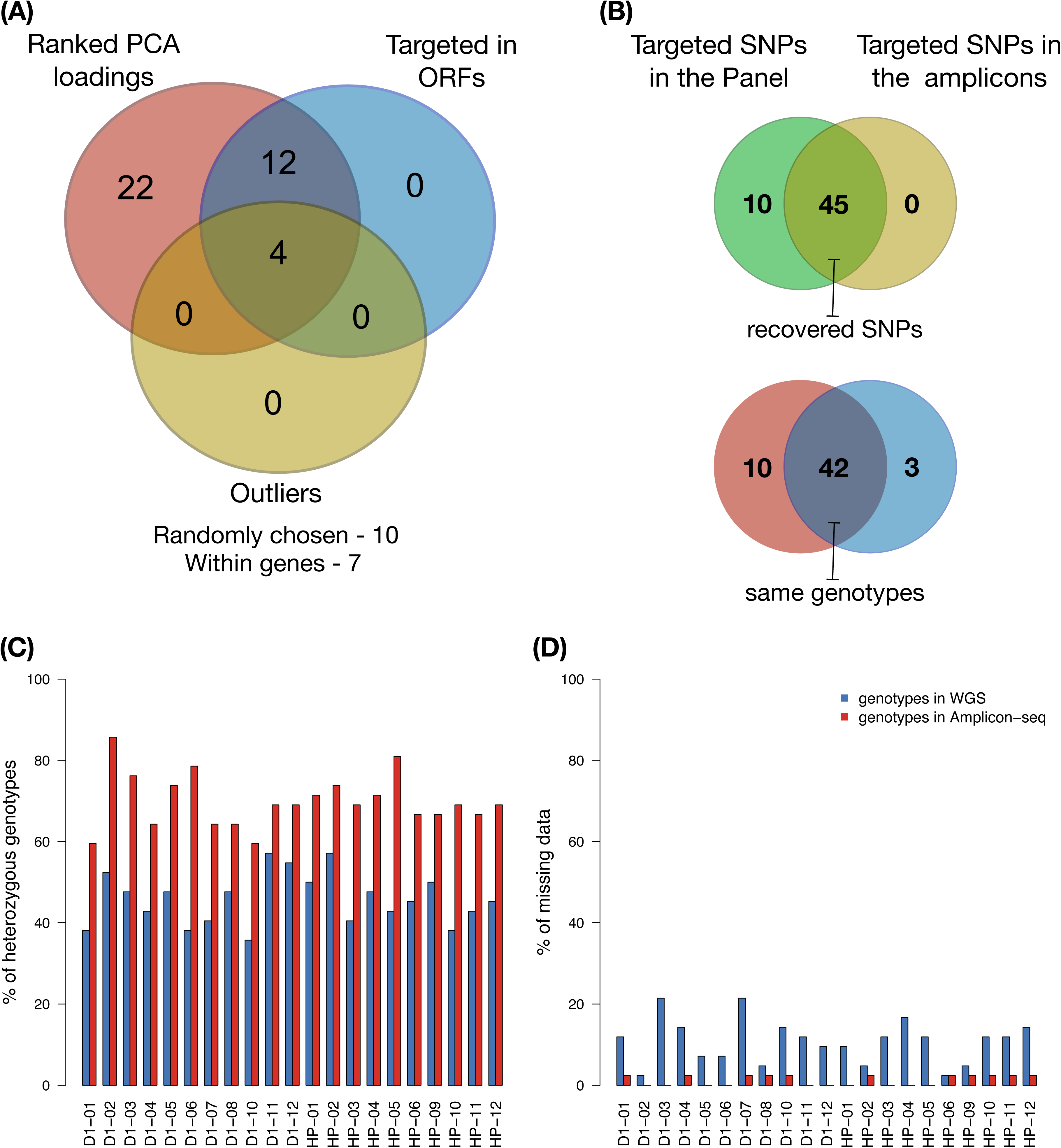
Summary of the performance of amplicon-sequencing for targeted SNPs. A) the type and number of SNPs included in the SNP panel. B) the number of targeted SNPs that were recovered with the amplicon-sequencing. C and D) improved coverage obtained and reduced missing data with the amplicon-sequencing.

Two out of 47 primer pairs did not uniquely align to their expected genomic region and were discarded (reads from paralogous genomic region can bias the genotype calls towards heterozygous calls, Hohenlohe *et al.* (2011); McKinney *et al.*(2016); Ravindran *et al.* (2018)). The alignment of predicted amplicon sequences to other amplicons sequences showed that three pairs of sequences had overlapping regions at one of the sequence ends. These overlaps were found in amplicons that came from adjacent regions of the genome. The partial alignment of reads to other regions can also cause an excess of heterozygous calls. In our pipeline we minimized the chance of partial alignment by imposing a stringent limit on read similarity allowed for read clustering. The partial alignment was removed using the amplicon sequence as the reference and a more stringent threshold for read alignment. As an example of the effectiveness of the mentioned steps, the targeted SNPs Bant_tg26 and Bant_tg27 were found in the overlapping regions, but the amount of heterozygous calls is proportional to the other targeted SNPs (Figure S8). For other primer sets, alignment of predicted amplicon sequences to the *B. antarctica* scaffolds showed the best hit was the expected scaffold and position.

### Validation of the panel for target sequencing enrichment

#### SNP calling for amplicon sequencing

FAA was 81% successful in recovering a targeted SNPs (45 out of 55 targeted SNPs). Seven targeted SNPs that were not recovered by FAA had adequate coverage but only had the reference allele, and three targeted SNPs (all in the *Buffy* gene) were not recovered (Figure S9B). We discarded an additional 3 SNPs (Bant_tgt10, Bant_tgt40, and Bant_tgt55) because they had different genotypes compared to the WGR SNPs, despite passing QC filters. Therefore, we used 42 out of 55 targeted SNPs to compare WGR and amplicon-sequencing to evaluate their ability to recover true genotypes (Figure 3B).

#### Validation of targeted SNPs and comparison between WGR and amplicon-seq SNPs

The average percentage of genotype similarities between sequencing protocols was 59.5%. This number was low because the high sequence depth of amplicon-sequencing (∼3000x) allowed us to reduce the number of homozygous calls being called as heterozygous (false positives) and heterozygous calls being called as homozygous (false negatives). The proportion of false positives and false negatives that were resolved by amplicon-sequencing were 6.2% and 22.2%, respectively. With amplicon-sequencing, we were able to confidently call new heterozygous genotypes that increased the proportion of within-individual heterozygous genotypes (Figure 3C). We were also able to reduce the amount of missing data within-individual and within-targeted SNPs (Figure 3D), although in one SNP it increased. When we compared the intra-locus estimates of H_E_, *π*, Tajima’s D and *θ*_W_ calculated for each targeted SNP we could see differences between the sequencing protocols, but the average estimates were concordant (except for the H_E_, Figure S10 and S11). We were also able to confirm genotypes of outliers discovered with WGR. The outlier with the highest posterior F_ST_ had the same genotype in both sequencing data, two outliers had the proportion of heterozygosity increased (Figure S12), and one was not polymorphic with the amplicon-sequencing data.

Amplicon-seq SNPs revealed the overall pattern of an excess of intermediate-frequency variants that was observed in the folded allele-frequency spectrum of WGR SNPs (Figure 1C). amplicon-seq SNP also showed a similar pattern of overall genetic differentiation between populations identified with PCA (Figure 1D). However, the global estimate of Weir & Cockerham’s F_ST_ for Amplicon-seq SNP was higher (0.083, 95% CI of 0.046 to 0.122) than the estimates obtained with WGR SNPs (0.028,95% CI of 0.022 to 0.032). Estimates of intra-locus summary statistics were similar, but estimates obtained with amplicon-seq SNPs showed lower variance for all summary statistics except for (Figure 4).

**Figure 4.**
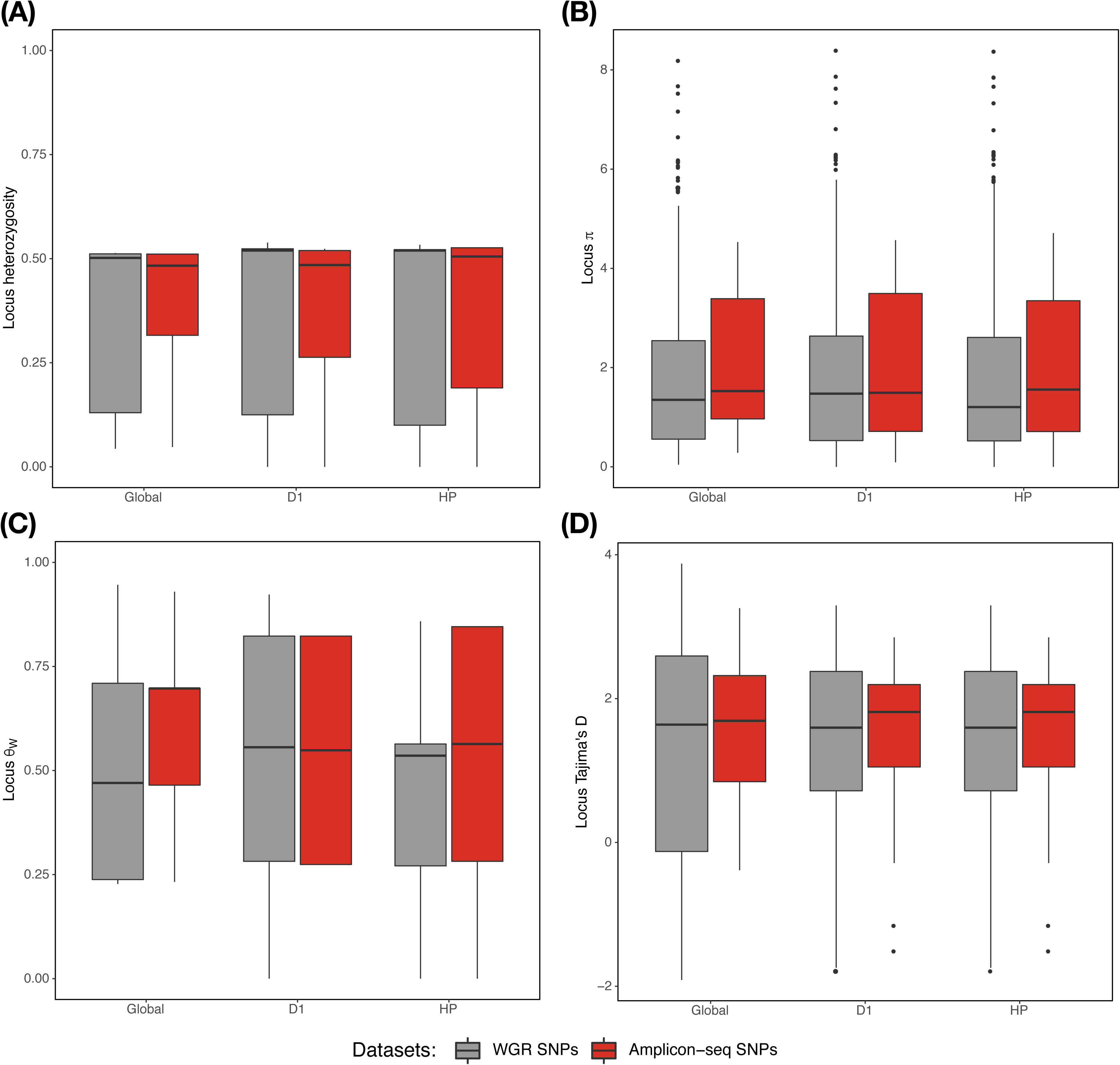
Comparison between summary statistics calculated with lcWGR and amplicon-sequencing SNPs. A) Expected heterozygosity - H_E_; B) nucleotide diversity – *π;* C) Watterson’s estimator - *θ*_W_; and D) Tajima’s D. In each plot the global and with-population summary statistic for each population were compared between sequencing protocol: WGR and Amplicon-sequencing. Dots represents the over-dispersed estimates.

## Discussion

For many populatio and conservation genetic studies with non-model organisms, it may be cost-prohibitive to generate high density WGR data sets. In this study, we show how the use of low-coverage, whole-genome re-sequencing (lcWGR) allows the identification of an informative set of SNP markers for rapid and reliable targeted enrichment genotyping. A cost-effective lcWGR was used to produce individual sequencing data to create a reduced set of informative SNPs for a SNP genotyping panel. Using the non-model dipteran, *B. antarctica*, the resultant SNP panel uncovered similar patterns of genetic diversity and population genetic structure as the lcWGR data set.

Coverage and quality of the lcWGR was low, limiting the identification of reliable variation. Using restrictive filters, we kept less than 1% of the identified variants, retaining SNPs in regions that had an average coverage higher than the expected 11X. The strict filtering came at a cost of discarding most rare to low-frequency variants (Fuentes-Pardo & Ruzzante, 2017) that may have shifted the folded-AFS spectrum. The low sample sizes for lcWGR also contributed to the AFS shift since the likelihood of a rare variant to be included was initially low. To minimize sample bias on the estimation of population genetic parameters (Albrechtsen, Nielsen & Nielsen, 2010; Fumagalli, 2013) and on the design of a reduced set of SNPs (Anderson, 2010; Ding *et al.*, 2011; Henriques *et al.*, 2018; Mariette *et al.*, 2002), a larger sample size would be required. However, in some cases with non-model organisms, large initial sample sizes may be challenging due to costs or unavailability of collections. While obtaining *B. antarctica* can be difficult from an isolated continent, we were fortunate to also have a complete genome for *B. antarctica*, which helped identify an informative set of markers. Regardless, focusing on medium-frequency to common SNPs increases the probability of recovering targeted SNPs and desired population genetic parameters for additional and distant populations.

The observed heterozygosity (H_O_) for the majority of WGR SNPs were higher compared to the expected heterozygosity (H_E_). The shift in folded-AFS also indicated accumulation of SNPs with a high proportion of heterozygous genotypes, indicating deviation from mutation-drift equilibrium. Both may be the consequence of the application of a hard filter on WGR SNPs, or alternatively may indicate two possible biological scenarios 1) strong bottlenecks events, and 2) a large population (with large N_e_) that underwent a recent admixture event with a close, but large and isolated, population. We can rule out an effect from QC filters since different sequencing protocols produced similar results (see below). The two possible biological scenarios, as well a combination of admixture and successive bottlenecks, are likely given the complex dynamics of seasonal freezing and thawing that is prevalent in Antarctica. However, with only 2 populations included in this study, inferences on demography would need additional sampling among several islands inhabited by *B. antarctica.*

In a scenario with strong stochastic changes in allele frequency and/or deviation from the island-model of migration with recent admixture (Bonhomme *et al.*, 2010; Whitlock & Lotterhos, 2015), we had limited power to identify outlier loci with F_ST_-based methods. Nonetheless, the combined methods identified one putative outlier locus that might be associated with stress adaptation in population D1, where all but one individual were heterozygous. The *B. antarctica* reference genome predicted that the alternative allele changed the amino acid from serine to arginine in a putative vitellogenin-A1 gene. Genotyping of other populations will allow us to see if differences in allele frequencies for the alternative allele exist.

The goal of our reduced SNP set was to recover and estimate summary statistics (H_E_, *π*, Tajima’s D, *θ*_W_) with some degree of similarity with WGR SNPs (similar average estimates and proportional variance range) and to rapidly assess natural populations. Despite differences in platforms, both produced relatively congruent results for the overall genetic diversity and population structure. It is interesting that we did not observe major discordance for the summary statistics among data types, as we obtained similar patterns for the folded-AFS, similar average estimates of H_E_, *π*, and e_W_, and similar individual assignments with PCA. We might attribute the ability of both WGR and Amplicon-seq SNPs to recover similar patterns of genetic diversity and population structure to the small genome of *B. antarctica.* Evolutionary events such as recent admixture and bottlenecks produce genome wide patterns on the genome; in small genomes their impact might be more extreme. In this case, the lcWGR data likely provided loci informative about overall patterns of the genome (*e.g.* diversity and demography). We had limited power to identify loci-specific features (*e.g.* selection), since bottlenecks and admixture could remove such signals. Nonetheless, for conservation genetic research, estimating genetic diversity and population structure is a more likely first step before identifying locus-specific adaptation.

Application of the reduced SNP strategy with the pipeline was shown to be effective for revealing major features underlying the evolutionary history of *B. antarctica*. We also observed strong clustering and differentiation among islands less than 8 kms apart, indicating some degree of isolation. These SNPs can be used more economically in multiple populations to better understand its current genetic diversity, describe global demography events, and predict any threats to extinction. For periodic genetic monitoring of this species, the ability to rapidly estimate genetic diversity is critical to identify issues that may be responding to drastic environmental changes across its wide range of inhabited Antarctic islands.

We also created a pipeline that automates the processing of Illumina’s short reads, produced by double-PCR amplification (that can be used with any amplicon sequencing protocol), and to generate full genotypes for individuals. With the amplicon-seq’s high coverage, this pipeline reduces false positives and negatives that impact the correct calling of heterozygous genotypes. The step-by-step guide for marker discovery and the pipeline designed for amplicon-sequencing can be used to isolate informative SNPs and rapidly genotype any non-model species. Amplicon-sequencing can not only speed-up the genotyping of endangered species but also facilitate the transition from conservation genetics to genomics (Meek & Larson, 2019; Taylor *et al.*, 2017). The pipeline is flexible enough for amplicon-sequencing with different degrees of amplicons and samples, and can also be used to prepare the data of low-coverage amplicon-sequencing experiments that incorporate some error (*e.g.* ANGSD, Korneliussen *et al.* (2014)). For those with less bioinformatic experience, our pipeline can rapidly provide input files needed for more user-friendly, non-command line, population genetic software.

## Conclusion

In summary, we show how amplicon-sequencing can be an alternative for population WGR when the goal is to acquire reliable genotypic information for many individuals and populations. When budget and bioinformatic experience limit the number of individuals to be sequenced with a decent sequence coverage, amplicon-sequencing offers a more affordable method compared to WGR or pool-seq. The process of isolating informative SNPs for targeted sequencing allows accurate recovery of fundamental estimates of genetic diversity and demographic patterns that are very similar to those generated by WGR. The guidelines presented here also include a pipeline developed to automate the processing of raw data produced by a combination of high throughput multiplex PCR and short-read sequencing into easily formatted genotyping input files. Our pipeline not only automates the genotyping but also reduces accumulation of sequencing errors, thereby increasing quality of genotyping calls. We believe this approach will be beneficial to many other non-model systems where questions concerning population and conservation genetic structure must be answered quickly to generate baseline data and predict future changes under climate change.

## Acknowledgements

We would like to thank M. Hernandez-Garcia at the MCIC for assistance with primer design and the FAA. Sarah Rudawsky helped with DNA extractions and quality assessment. Funding was provided by NSF Division of Polar Programs (35000404): PLR-1341393.

## Data availability

WGR and amplicon-sequencing data are available at NCBI BioProject accession numbers: PRJNA565001 and PRJNA565153.The sequence of each primer pair, the reference amplicons (fasta file) and R scripts are available at https://github.com/vitorpavinato/belgicaampliconseq. The pipeline to process amplicon-sequencing raw data – PypeAmplicon – is available at zendo (doi:10.5281/zenodo.1490421) and on GitHub (https://github.com/vitorpavinato/PypeAmplicon).

## Author Contributions

A.P.M., D.L.D., T.M., and V.A.C.P. designed the research; V.A.C.P analyzed the WGS and the Amplicon-sequencing data; V.A.C.P and D.S developed the reduced SNP panel; V.A.C.P. conduct laboratory work on the Amplicon-sequencing; A.P.M., and T.M. provided support for lab work; S.W and V.A.C.P developed the Amplicon-sequencing pipeline; S.W provided bioinformatics consultancy; V.A.C.P., A.P.M., and D.L.D. wrote the manuscript.

## Notes

https://github.com/vitorpavinato/belgicaampliconseq

https://github.com/vitorpavinato/PypeAmplicon

## References

Albrechtsen, A., Nielsen, F. C. & Nielsen, R. (2010). Ascertainment biases in SNP Chips affect measures of population divergence. Molecular Biology and Evolution, 27(11), 2534–2547. doi:10.1093/molbev/msq148

Allendorf, F. W. (2017). Genetics and the conservation of natural populations: allozymes to genomes. Molecular Ecology, 26(2), 420–430. doi:10.1111/mec.13948

Allegrucci, G., Carchini, G., Todisco, V., Convey, P., Sbordoni, V.. (2006). A molecular phylogeny of antarctic chironomidae and its implications for biogeographical history. Polar Biology 29(4):320–326. doi:10.1007/s00300-005-0056-7

Altschul, S. F., Gish, W., Miller, W., Myers, E. W. & Lipman, D. J. (1990). Basic local alignment search tool. Journal of Molecular Biology, 215(3), 403–410. doi:10.1016/s0022-2836(05)80360-2

Anderson, E. C. (2010). Assessing the power of informative subsets of loci for population assignment: standard methods are upwardly biased. Molecular Ecology Resources, 10(4), 701–710. doi:10.1111/j.1755-0998.2010.02846.x

Baird, N.A., Etter, P. D., Atwood, T.S, Currey, M.C., Shiver, A. L., Lewis, Z. A., Selker, E. U., … Johnson, E. A. (2008). Rapid SNP discovery and genetic mapping using sequenced RAD markers. PLoS Genetics, 3(10), e3376–7. Doi:10.1371/journal.pone.0003376

Bolger, A. M., Lohse, M. & Usadel, B. (2014). Trimmomatic: a flexible trimmer for Illumina sequence data. Bioinformatics, 30(15), 2114–2120. doi:10.1093/bioinformatics/btu170

Bonhomme, M., Chevalet, C., Servin, B., Boitard, S., Abdallah, J., Blott, S. & SanCristobal, M. (2010). Detecting selection in population trees: the Lewontin and Krakauer test extended. Genetics, 186(1), 241–262. doi:10.1534/genetics.110.117275

Bowden, R., MacFie, T. S., Myers, S., Hellenthal, G., Nerrienet, E., Bontrop, R. E., Freeman, C., … Mundy N. I. (2012). Genomic tools for evolution and conservation in the Chimpanzee: *Pan troglodytes ellioti* is a genetically distinct population. PLoS Genetics, 8(3), e1002504. doi:10.1371/journal.pgen.1002504

Brown, A. P., McGowan, K. L., Schwarzkopf, E. J., Greenway, R., Rodriguez, L. A., Tobler, M. & Kelley, J. L. (2019). Local ancestry analysis reveals genomic convergence in extremophile fishes. Philosophical Transactions of the Royal Society B: Biological Sciences, 374(1777), 20180240. doi:10.1098/rstb.2018.0240

Campbell, N. R., Harmon, S. A. & Narum, S. R. (2014). Genotyping-in-Thousands by sequencing (GT-seq): a cost effective SNP genotyping method based on custom amplicon sequencing. Molecular Ecology Resources, 15(4), 855–867. doi:10.1111/1755-0998.12357

Convey, P. & Block, W. (1996). Antartic Diptera: ecology, physiology and distribution. European Journal of Entomology, 93 (1), 1–13.

Danecek, P., Auton, A., Abecasis, G., Albers, C. A., Banks, E., DePristo, M. A., Handsaker, R. E., … Durbin, R. (2011). The variant call format and VCFtools. Bioinformatics, 27(15), 2156–2158. doi:10.1093/bioinformatics/btr330

Danial, N. N. & Korsmeyer, S. J. (2004). Cell death: critical control points. Cell, 116(2), 205–219. doi:10.1016/s0092-8674(04)00046-7

Ding, L., Wiener, H., Abebe, T., Altaye, M., Go, R. C. P., Kercsmar, C., Grabowski, G., … Baye, T. M. (2011). Comparison of measures of marker informativeness for ancestry and admixture mapping. BMC Genomics, 12(1), 622. doi:10.1186/1471-2164-12-622

Edgar, R. C. (2010). Search and clustering orders of magnitude faster than BLAST. Bioinformatics, 26(19), 2460–2461. doi:10.1093/bioinformatics/btq461

Ekblom, R. & Galindo, J. (2010). Applications of next generation sequencing in molecular ecology of non-model organisms. Heredity, 107(1), 1–15. doi:10.1038/hdy.2010.152

Fischer, M. C., Rellstab, C., Leuzinger, M., Roumet, M., Gugerli, F., Shimizu, K. K., … Widmer, A. (2017). Estimating genomic diversity and population differentiation - an empirical comparison of microsatellite and SNP variation in *Arabidopsis halleri*. BMC Genomics, 18(1), 69. doi:10.1186/s12864-016-3459-7

Foll, M. & Gaggiotti, O. (2008). A genome-scan method to identify selected loci appropriate for both dominant and codominant markers: a bayesian perspective. Genetics, 180(2), 977–993. doi:10.1534/genetics.108.092221

Fuentes-Pardo, A. P. & Ruzzante, D. E. (2017). Whole-genome sequencing approaches for conservation biology: advantages, limitations and practical recommendations. Molecular Ecology, 26(20), 5369–5406. doi:10.1111/mec.14264

Fumagalli, M. (2013). Assessing the effect of sequencing depth and sample size in population genetics inferences. PLoS ONE, 8(11), e79667. doi:10.1371/journal.pone.0079667

Garrison, E. & Marth, G. (2012). Haplotype-based variant detection from short-read sequencing. org/abs/1207.3907

Goudet, J. (2005). hierfstat, a package for r to compute and test hierarchical F-statistics. Molecular Ecology Notes, 5(1), 184–186. doi:10.1111/j.1471-8286.2004.00828.x

Hayward, S. A. L., Rinehart, J. P., Sandro, L. H., Lee, R. E. & Denlinger, D. L. (2007). Slow dehydration promotes desiccation and freeze tolerance in the Antarctic midge *Belgica antarctica*. Journal of Experimental Biology, 210(5), 836–844. doi:10.1242/jeb.02714

Henriques, D., Parejo, M., Vignal, A., Wragg, D., Wallberg, A., Webster, M. T. & Pinto, M. A. (2018). Developing reduced SNP assays from whole-genome sequence data to estimate introgression in an organism with complex genetic patterns, the Iberian honeybee (*{A}pis mellifera iberiensis*). Evolutionary Applications, 11(8), 1270–1282. doi:10.1111/eva.12623

Hill, R. J., Billas, I. M. L., Bonneton, F., Graham, L. D. & Lawrence, M. C. (2013). Ecdysone receptors: from the Ashburner model to structural biology. Annual Review of Entomology, 58(1), 251–271. doi:10.1146/annurev-ento-120811-153610

Hohenlohe, P. A., Amish, S. J., Catchen, J. M., Allendorf, F. W. & Luikart, G. (2011). Next-generation RAD sequencing identifies thousands of SNPs for assessing hybridization between rainbow and westslope cutthroat trout. Molecular Ecology Resources, 11, 117–122. doi:10.1111/j.1755-0998.2010.02967.x

Hulsegge, B., Calus, M. P. L., Windig, J. J., Hoving-Bolink, A. H., Eijndhoven, M. H. T. M.-v. & Hiemstra, S. J. (2013). Selection of SNP from 50K and 777K arrays to predict breed of origin in cattle. Journal of Animal Science, 91(11), 5128–5134. doi:10.2527/jas.2013-6678

Jombart, T. (2008). adegenet: a R package for the multivariate analysis of genetic markers. Bioinformatics, 24(11), 1403–1405. doi:10.1093/bioinformatics/btn129

Kelley, J. L., Peyton, J. T., Fiston-Lavier, A.-S., Teets, N. M., Yee, M.-C., Johnston, J. S., Bustamante, C. D., … Denlinger, D. L. (2014). Compact genome of the Antarctic midge is likely an adaptation to an extreme environment. Nature Communications, 5(1), 4611. doi:10.1038/ncomms5611

Korneliussen, T. S., Albrechtsen, A. & Nielsen, R. (2014). ANGSD: analysis of next generation sequencing data. BMC Bioinformatics, 15(1), 356. doi:10.1186/s12859-014-0356-4

Langmead, B. & Salzberg, S. L. (2012). Fast gapped-read alignment with Bowtie 2. Nature Methods, 9(4), 357–359. doi:10.1038/nmeth.1923

Larson, W. A., Dann, T. H., Limborg, M. T., McKinney, G. J., Seeb, J. E. & Seeb, L. W. (2019). Parallel signatures of selection at genomic islands of divergence and the major histocompatibility complex in ecotypes of sockeye salmon across Alaska. Molecular Ecology, 28(9), 2254–2271,. doi:10.1111/mec.15082

Lee, Y. S., Nakahara, K., Pham, J. W., Kim, K., He, Z., Sontheimer, E. J. & Carthew, R. W. (2004). Distinct roles for Drosophila Dicer-1 and Dicer-2 in the siRNA/miRNA silencing pathways. Cell, 117(1), 69–81. doi:10.1016/s0092-8674(04)00261-2

Li, H., Handsaker, B., Wysoker, A., Fennell, T., Ruan, J., Homer, N., Marth, G., … 1000 Genome Project Data Processing Subgroup (2009). The sequence alignment/map format and SAMtools. Bioinformatics, 25(16), 2078–2079. doi:10.1093/bioinformatics/btp352

Mace, G. M. (2004). The role of taxonomy in species conservation. Philosophical Transactions of the Royal Society of London. Series B: Biological Sciences, 359(1444), 711–719. doi:10.1098/rstb.2003.1454

Mariette, S., Corre, V. L., Austerlitz, F. & Kremer, A. (2002). Sampling within the genome for measuring within-population diversity: trade-offs between markers. Molecular Ecology, 11(7), 1145–1156. doi:10.1046/j.1365-294x.2002.01519.x

McKenna, A., Hanna, M., Banks, E., Sivachenko, A., Cibulskis, K., Kernytsky, A., Garimella, K., … DePristo M. A. (2010). The genome analysis toolkit: a MapReduce framework for analyzing next-generation DNA sequencing data. Genome Research, 20(9), 1297–1303. doi:10.1101/gr.107524.110

McKinney, G. J., Waples, R. K., Seeb, L. W. & Seeb, J. E. (2016). Paralogs are revealed by proportion of heterozygotes and deviations in read ratios in genotyping-by-sequencing data from natural populations. Molecular Ecology Resources, 17(4), 656–669. doi:10.1111/1755-0998.12613

Meek, M. H. & Larson, W. A. (2019). The future is now: amplicon sequencing and sequence capture usher in the conservation genomics era. Molecular Ecology Resources, 19(4), 795–803. doi:10.1111/1755-0998.12998

Milano, I., Babbucci, M., Cariani, A., Atanassova, M., Bekkevold, D., Carvalho, G.R, Espiñeira M., … Bargelloni, L. (2013). Outlier SNP markers reveal fine-scale genetic structuring across European hake populations (*Merluccius merluccius*). Molecular Ecology, 23(1), 118–135. doi:10.1111/mec.12568

Mita, S. D. & Siol, M. (2012). EggLib: processing, analysis and simulation tools for population genetics and genomics. BMC Genetics, 13(1), 27. doi:10.1186/1471-2156-13-27

Paradis, E. (2010). pegas: an R package for population genetics with an integrated-modular approach. Bioinformatics, 26(3), 419–420. doi:10.1093/bioinformatics/btp696

Peterson, B. K., Weber, J. N., Kay, E. H., Fisher, H. S. & Hoekstra, H. E. (2012). Double digest RADseq: an inexpensive method for de novo SNP discovery and genotyping in model and non-model species. PLoS ONE, 7(5), e37135. doi:10.1371/journal.pone.0037135

Prado-Martinez, J., Sudmant, P.H., Kidd, J.M., Li, H., Kelley, J.L., Lorente-Galdos, B., Veeramah, K.R., … Marques-Bonet T., (2013). Great ape genetic diversity and population history. Nature, 499(7459), 471–475. doi:10.1038/nature12228

Quinn, L. (2003). Buffy, a Drosophila Bcl-2 protein, has anti-apoptotic and cell cycle inhibitory functions. The EMBO Journal, 22(14), 3568–3579. doi:10.1093/emboj/cdg355

Ravindran, P. N., Bentzen, P., Bradbury, I. R. & Beiko, R. G. (2018). PMERGE: Computational filtering of paralogous sequences from RAD-seq data. Ecology and Evolution, 8(14), 7002–7013. doi:10.1002/ece3.4219

Rinehart, J. P., Hayward, S. A. L., Elnitsky, M. A., Sandro, L. H., Lee, R. E. & Denlinger, D. L. (2006). Continuous up-regulation of heat shock proteins in larvae, but not adults, of a polar insect. Proceedings of the National Academy of Sciences, 103(38), 14223–14227. doi:10.1073/pnas.0606840103

Supple, M. A. & Shapiro, B. (2018). Conservation of biodiversity in the genomics era. Genome Biology, 19(1), 131. doi:10.1186/s13059-018-1520-3

Tajima, F. (1989). Statistical method for testing the neutral mutation hypothesis by DNA polymorphism. Genetics, 123(3), 585–595.

Taylor, H. R., Dussex, N. & van Heezik, Y. (2017). Bridging the conservation genetics gap by identifying barriers to implementation for conservation practitioners. Global Ecology and Conservation, 10, 231–242. doi:10.1016/j.gecco.2017.04.001

Teets, N. M., Kawarasaki, Y., Lee, R. E. & Denlinger, D. L. (2012b). Expression of genes involved in energy mobilization and osmoprotectant synthesis during thermal and dehydration stress in the Antarctic midge, *Belgica antarctica*. Journal of Comparative Physiology B, 183(2), 189–201. doi:10.1007/s00360-012-0707-2

Teets, N. M., Peyton, J. T., Colinet, H., Renault, D., Kelley, J. L., Kawarasaki, Y., …, Denlinger D. L. (2012a). Gene expression changes governing extreme dehydration tolerance in an Antarctic insect. Proceedings of the National Academy of Sciences, 109(50), 20744–20749. doi:10.1073/pnas.1218661109

Watterson, G. A. (1975). On the number of segregating sites in genetical models without recombination. Theoretical Population Biology, 7(2), 256–276.

Whitlock, M. C. & Lotterhos, K. E. (2015). Reliable detection of loci responsible for local adaptation: inference of a null model through trimming the distribution of FST. The American Naturalist, 186(S1), S24–S36. doi:10.1086/682949

Wilkinson, S., Wiener, P., Archibald, A. L., Law, A., Schnabel, R. D., McKay, S.D., … Ogden R. (2011). Evaluation of approaches for identifying population informative markers from high density SNP Chips. BMC Genetics, 12(1), 45. doi:10.1186/1471-2156-12-45

Wijeratne, S. & Pavinato, V. A. C. (2018). PypeAmplicon v1.0: Python pipeline for analysis of amplicon data. Zenodo. doi:10.5281/zenodo.1490421

Yang, S., Fresnedo-Ramírez, J., Wang, M., Cote, L., Schweitzer, P., Barba, P., Takacs, E.M., … Sun, Q. (2016). A next-generation marker genotyping platform (AmpSeq) in heterozygous crops: a case study for marker-assisted selection in grapevine. Horticulture Research, 3(1), 16002. doi:10.1038/hortres.2016.2m,

