## Supplementary Materials for "Leveraging targeted sequencing for non-model species: a step-by-step guide to obtain a reduced SNP set and a pipeline to automate data processing in the Antarctic Midge, *Belgica antarctica*"

**Table S1. Outlier loci detected with BAYESCAN and OUTFLANK**

**Table S2. Corrected genotypes with amplicon sequencing.**

**Figure S1. Collection sites of samples used for the marker discovery and amplicon-sequencing SNP targeted validation.**

**Figure S2. Reference-based pipeline for de novo variant discovery of lcWGR.** Blue represent data and grey the programs used in each step of the marker discover pipeline for whole genome sequencing data.

**FigureS3. Population genetics-based filtering of informative SNPs.** 1) The loadings of a PCA. All WGR SNPs were ranked and SNPs associated with the highest values were selected. 2) SNPs that were in H-W proportion based on Fisher's exact test were retained 3) SNPs with only heterozygous genotypes or surrounded by SNPs with all heterozygous genotypes were removed. SNPs within or near a cluster of SNPs were also removed. 4) SNP detected as outliers with BAYESCAN and OUTFLANK. 5) SNPs within targeted genes. We also included random SNPs (that passed the above mentioned steps except the PCA loadings ranking).

**Figure S4. Analytical workflow for analyzing reads from amplicon sequencing.** Orange represents steps to process the amplicon data (in blue are the data produced for each step). Paired-end reads were first merged before quality control steps. Merged reads were clustered by similarities and orphaned reads or clusters with few reads were discarded. Only clusters of reads that mapped to the reference amplicon sequencing were retained. Reads were then aligned to the reference amplicon to produce BAM files for each amplicon/individual. The last steps merged BAM (amplicon/individual) and the SNP calling.

**Figure S5. Average coverage for the lcWGR protocol.** A) the average coverage estimates for the three steps of alignment-based analysis. B) histogram showing the distribution of mean sequencing depth for the filtered SNPs.

**Figure S6. Fisher's exact test for H-W proportions on lcWGR SNPs.** A, B and C) quantile-quantile plots of p-values calculated for each locus with Fisher's exact test. D) the proportion of loci with departure from H-W proportion without multiple test corrections.

**Figure S7. LD decay graph of lcWGR SNPs.**

**Figure S8. Comparison between the proportion of heterozygous individuals for each targeted SNPs for lcWGR and amplicon-sequencing protocols.**

**Figure S9. Average depth for targeted SNPs.** A) show the average depth of each targeted SNPs obtained with lcWGR. B) shows the average depth obtained with the amplicon-sequencing.

**Figure S10 and S11. Comparison between intra-locus summary statistics of targeted SNPs.** Each locus estimate for the summary statistics ($H_E$, $\pi$, $\Theta_W$ and Tajima's D) was compared between sequencing protocols.

**Figure S12. Concordance between the genotypes of outlier loci.** To illustrate the applicability of amplicon-sequencing to validate outliers detected with lcWGSR, this graph compares the number of genotypes for each possible genotype of bi-allelic SNPs (Ref/Ref, Ref/Alt, Alt/Alt), obtained with the WGR and amplicon-sequencing.

| Scaffold | Position | SNP | Targeted name | Alleles (Ref,Alt) | Posterior FST | Pi | | Ensembl Variant Effect Predictor – VEP | | | Homology | | |
|---|---|---|---|---|---|---|---|---|---|---|---|---|---|
| | | | | | | D1 | HP | Consequence | Impact | Gene1 | Protein2 | Gene | GO |
| JPYR01001350 | 7449 | 490 | - | A, G | 0.030 | 1.348 | 2.012 | Synonymous variant | Low | IU25_12415 | Uncharacterized protein | - | - |
| JPYR01001350 | 7518 | 492 | - | G, A | 0.046 | 1.348 | 2.012 | Synonymous variant | Low | IU25_12415 | Uncharacterized protein | - | - |
| JPYR01001535 | 416 | 611 | Bant_tgt17 | G, T | 0.061 | 0.519 | 0.000 | Missense variant | Moderate | IU25_12621 | CLUMA_CG006889, isoform A | Putative Vitellogenin-A1 | Lipid transporter activity |
| JPYR01001535 | 1491 | 612 | Bant_tgt18 | G, T | 0.028 | 0.503 | 0.000 | Missense variant | Moderate | IU25_12621 | CLUMA_CG006889, isoform A | Putative Vitellogenin-A1 | Lipid transporter activity |
| JPYR01001792 | 2458 | 688 | Bant_tgt36 | T, C | 0.024 | 0.363 | 2.382 | Synonymous variant | Low | IU25_12829 | Uncharacterized protein | - | - |
| JPYR01001792 | 2713 | 693 | Bant_tgt37 | T, G | 0.024 | 0.289 | 2.021 | Missense variant | Moderate | IU25_12829 | Uncharacterized protein | - | - |
| JPYR01002934 | 360 | 1105 | - | G, A | 0.028 | 1.466 | 1.155 | Intergenic variant | Modifier | - | - | - | - |
| JPYR01004124 | 115 | 1213 | - | C, T | 0.030 | 0.523 | 0.100 | Intergenic variant | Modifier | - | - | - | - |

| | | Targeted SNPs | | | |
|---|---|---|---|---|---|
| | | RR | RA | AA | NA |
| Amplicon-seq SNPs | RR | **158** | 47 | 0 | 19 |
| | RA | 176 | **347** | 19 | 75 |
| | AA | 1 | 8 | **20** | 1 |
| | NA | 9 | 2 | 0 | **0** |

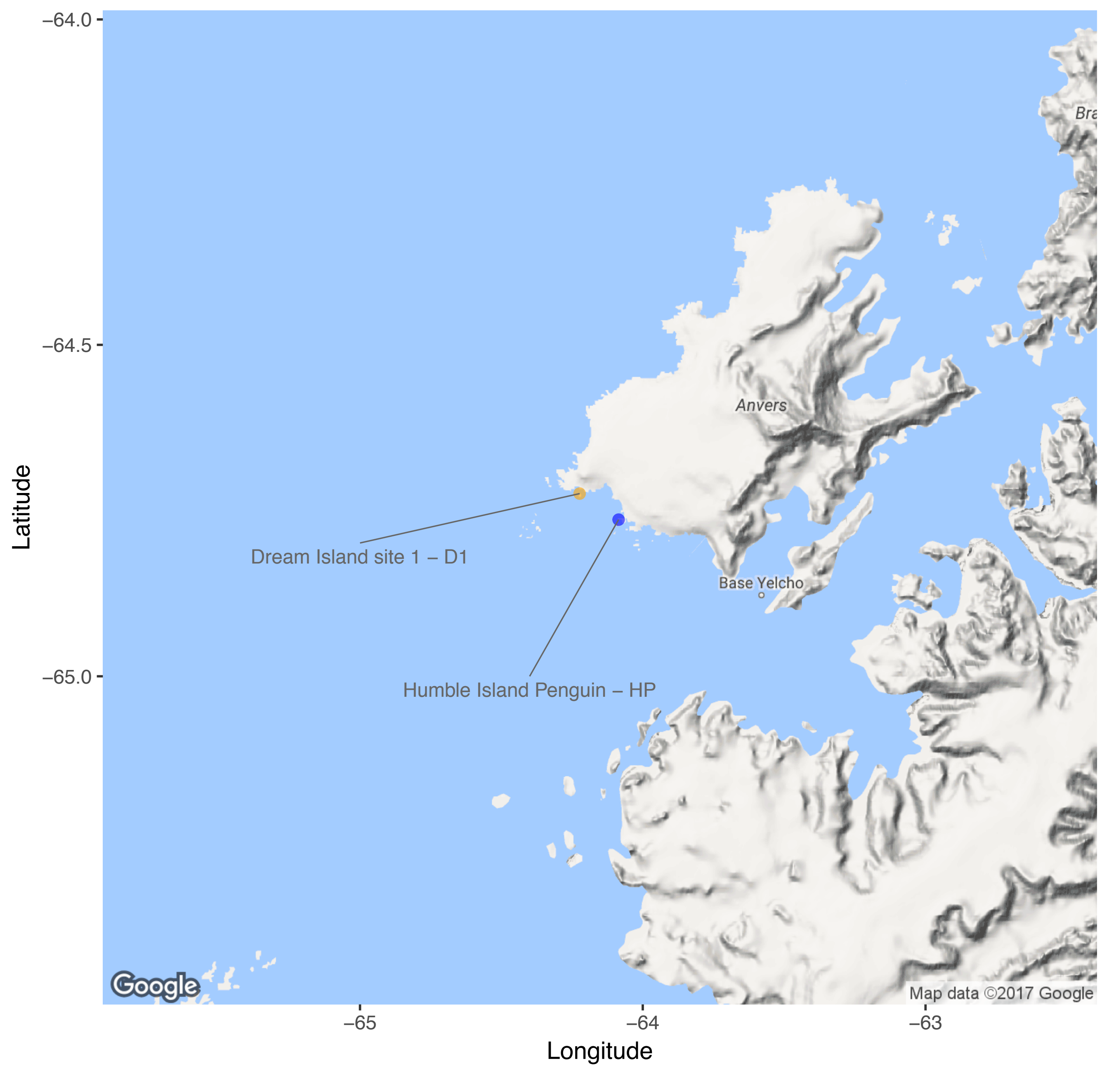

−64.0
−64.5
−65.0
Latitude
−65
−64
−63
Longitude
Bra
Anvers
Base Yelcho
Dream Island site 1 – D1
Humble Island Penguin – HP
Google
Map data ©2017 Google

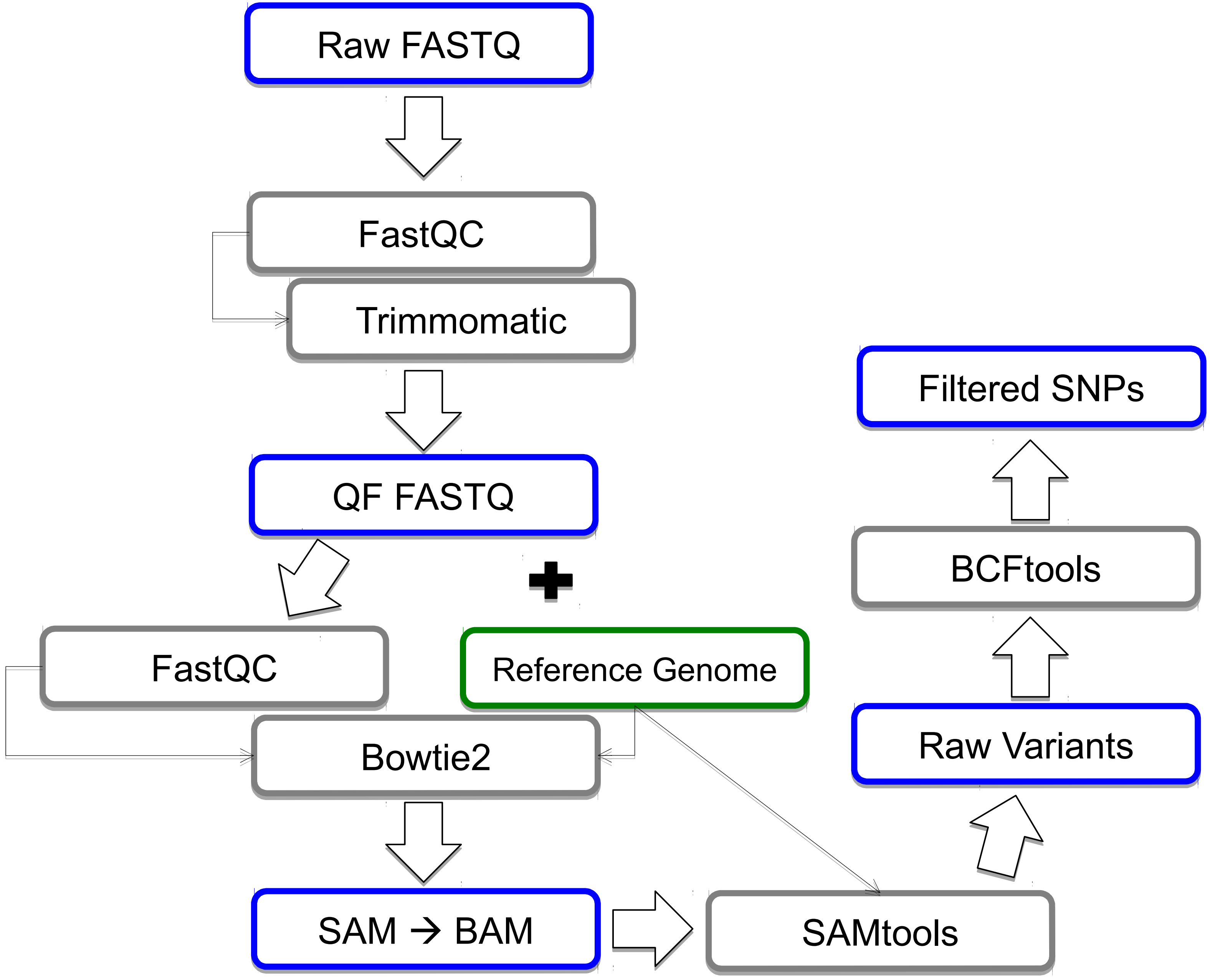

Raw FASTQ
FastQC
Trimmomatic
QF FASTQ
FastQC
Reference Genome
Bowtie2
SAM → BAM
SAMtools
Raw Variants
BCFtools
Filtered SNPs

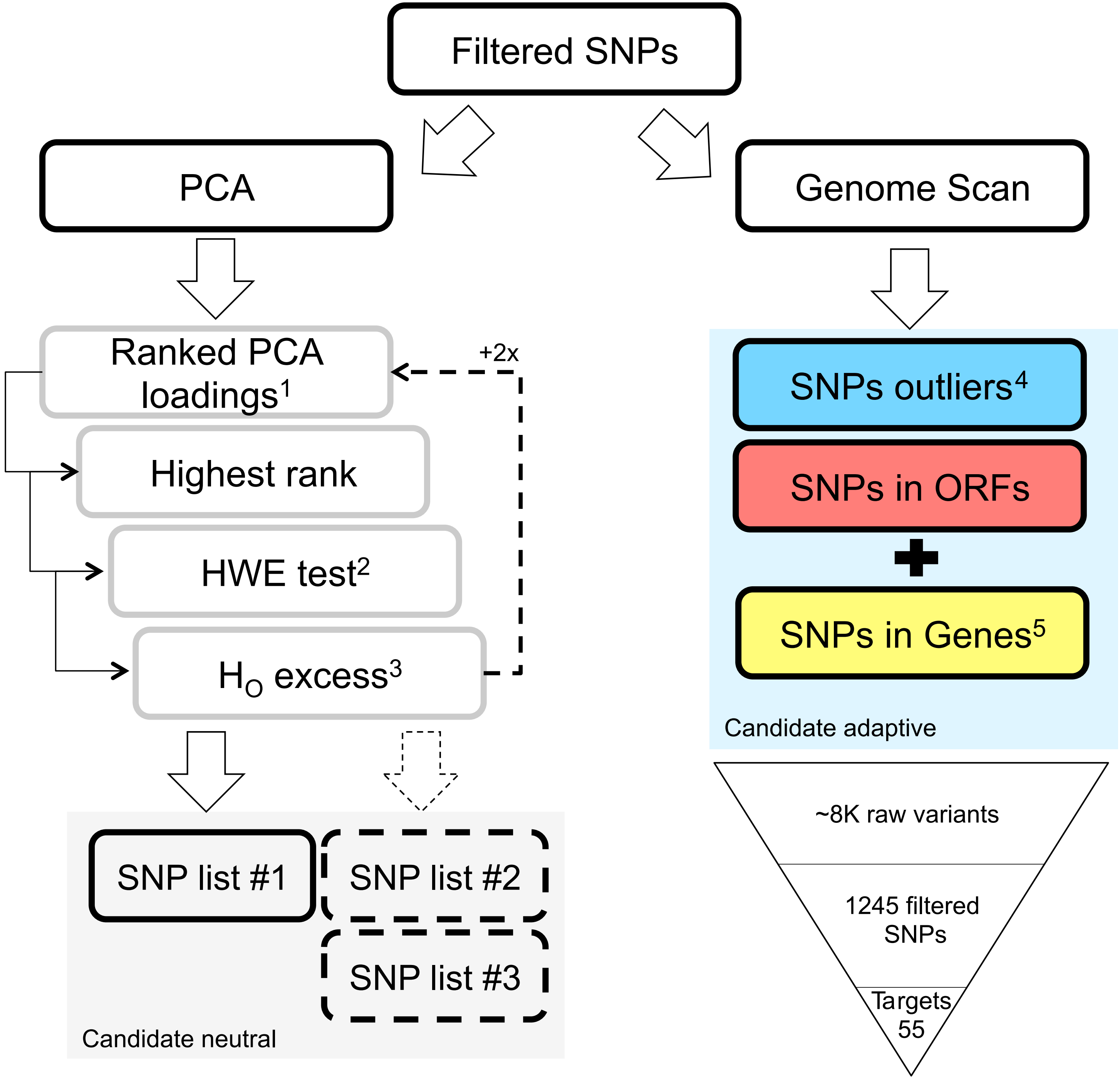
Filtered SNPs
PCA
Genome Scan
Ranked PCA loadings1
+2x
Highest rank
HWE test2
HO excess3
SNP list #1
SNP list #2
SNP list #3
Candidate neutral
SNPs outliers4
SNPs in ORFs
SNPs in Genes5
Candidate adaptive
~8K raw variants
1245 filtered SNPs
Targets 55

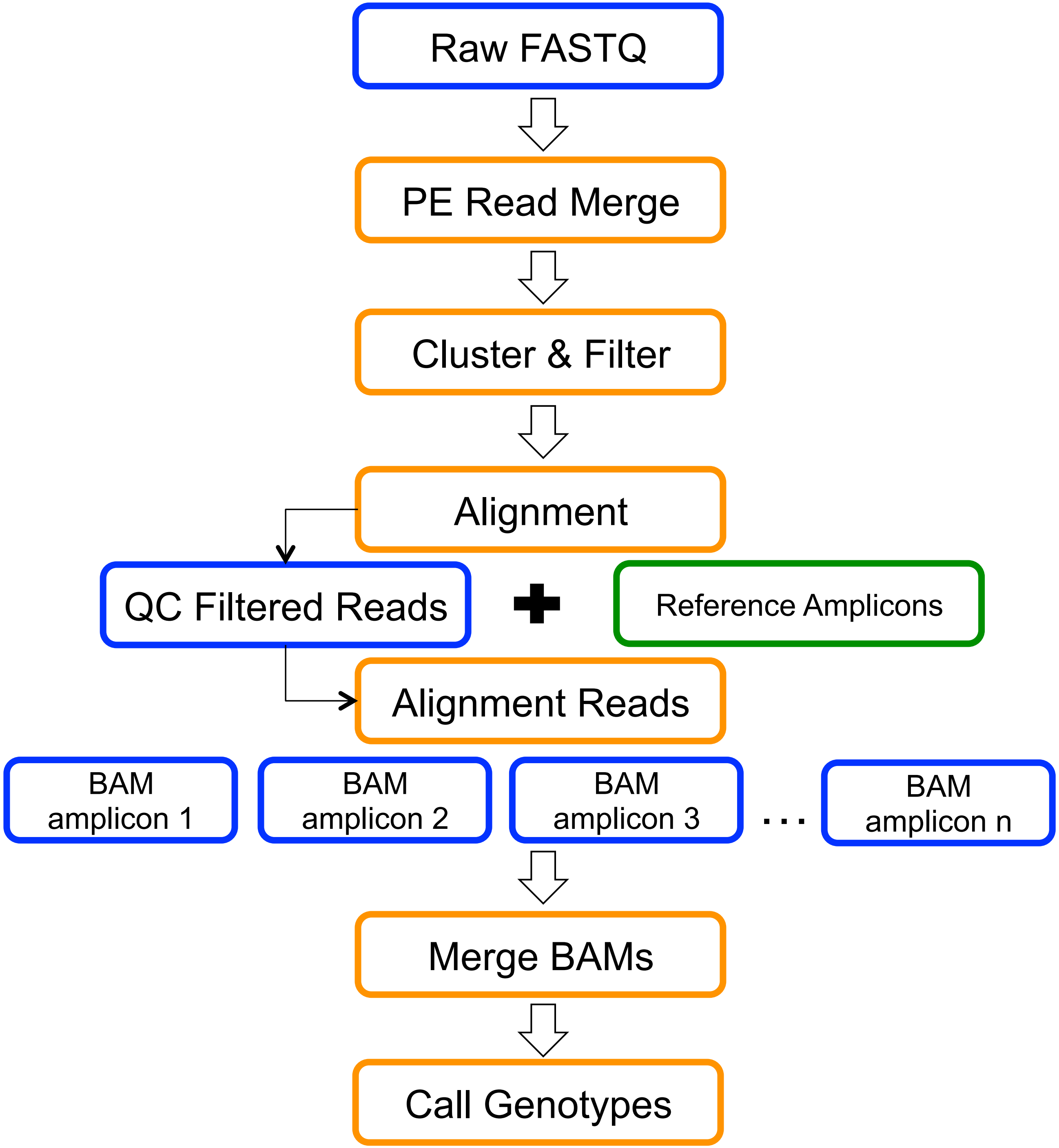
Raw FASTQ
PE Read Merge
Cluster & Filter
Alignment
QC Filtered Reads
Reference Amplicons
Alignment Reads
BAM amplicon 1
BAM amplicon 2
BAM amplicon 3
. . .
BAM amplicon n
Merge BAMs
Call Genotypes

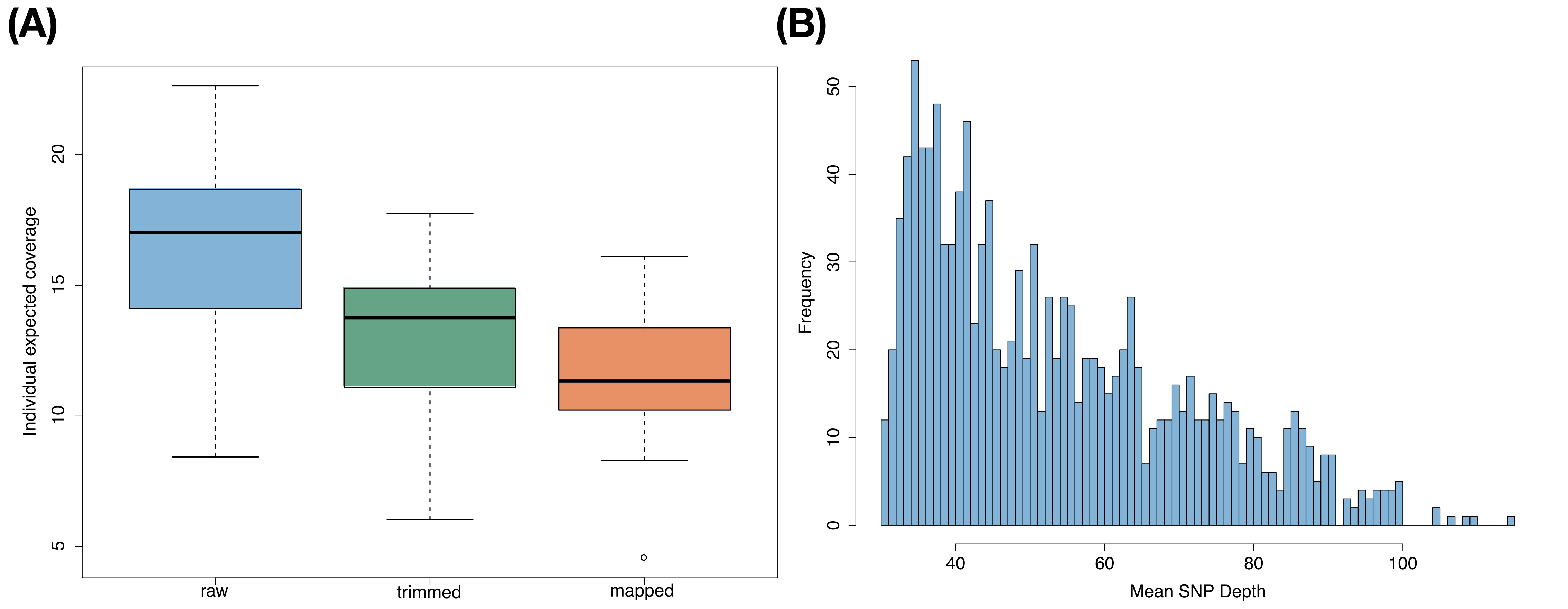

(A)
Individual expected coverage
5
10
15
20
raw
trimmed
mapped
(B)
Frequency
0
10
20
30
40
50
40
60
80
100
Mean SNP Depth

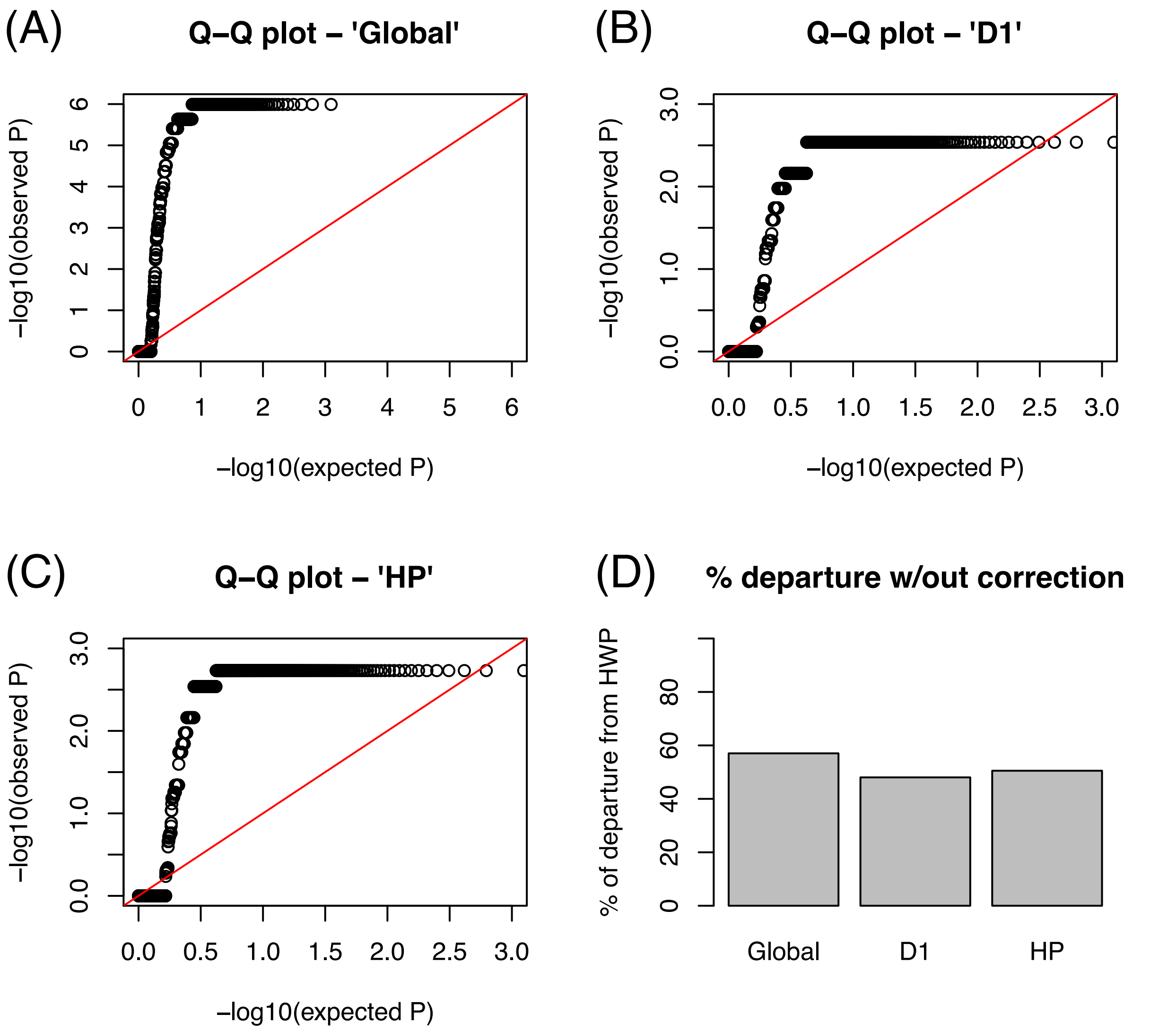
(A)
Q–Q plot – 'Global'
−log10(observed P)
−log10(expected P)
0
1
2
3
4
5
6
(B)
Q–Q plot – 'D1'
−log10(observed P)
−log10(expected P)
0.0
0.5
1.0
1.5
2.0
2.5
3.0
(C)
Q–Q plot – 'HP'
−log10(observed P)
−log10(expected P)
0.0
0.5
1.0
1.5
2.0
2.5
3.0
(D)
% departure w/out correction
% of departure from HWP
0
20
40
60
80
Global
D1
HP

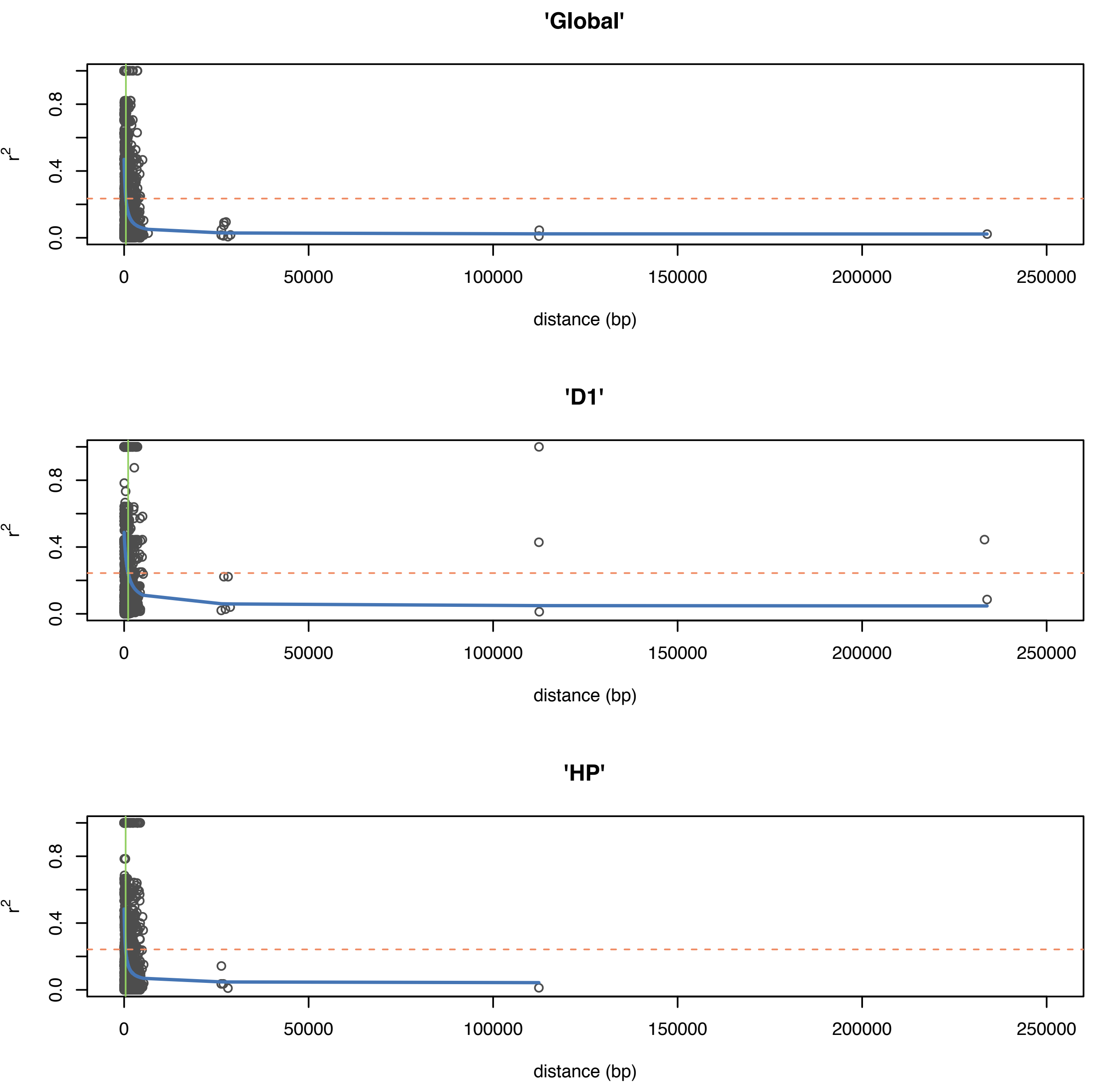

'Global'
'D1'
'HP'
r2
0.0
0.4
0.8
0
50000
100000
150000
200000
250000
distance (bp)

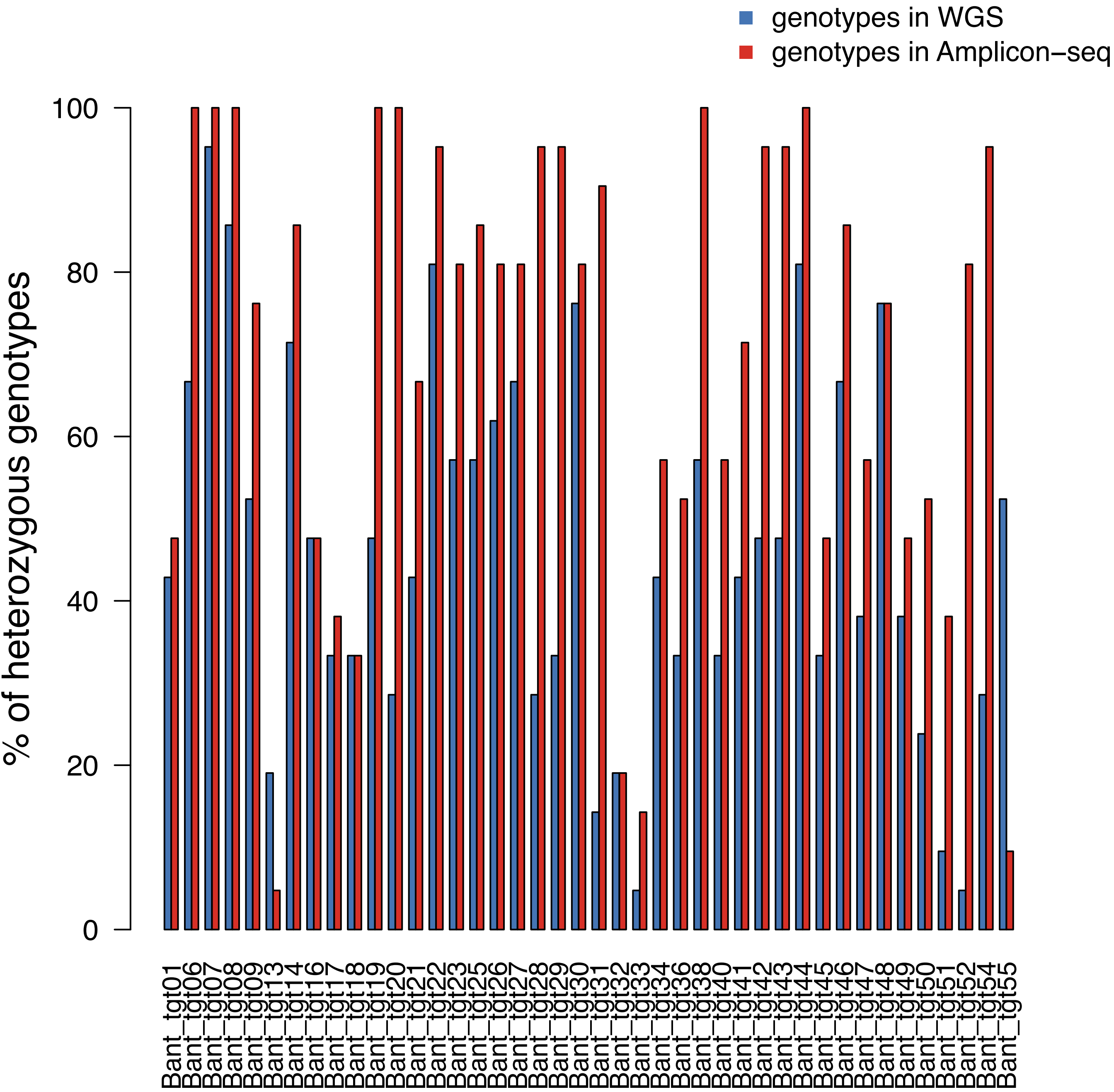

genotypes in WGS
genotypes in Amplicon−seq
% of heterozygous genotypes
0
20
40
60
80
100
Bant_tgt01
Bant_tgt06
Bant_tgt07
Bant_tgt08
Bant_tgt09
Bant_tgt13
Bant_tgt14
Bant_tgt16
Bant_tgt17
Bant_tgt18
Bant_tgt19
Bant_tgt20
Bant_tgt21
Bant_tgt22
Bant_tgt23
Bant_tgt25
Bant_tgt26
Bant_tgt27
Bant_tgt28
Bant_tgt29
Bant_tgt30
Bant_tgt31
Bant_tgt32
Bant_tgt33
Bant_tgt34
Bant_tgt36
Bant_tgt38
Bant_tgt40
Bant_tgt41
Bant_tgt42
Bant_tgt43
Bant_tgt44
Bant_tgt45
Bant_tgt46
Bant_tgt47
Bant_tgt48
Bant_tgt49
Bant_tgt50
Bant_tgt51
Bant_tgt52
Bant_tgt54
Bant_tgt55

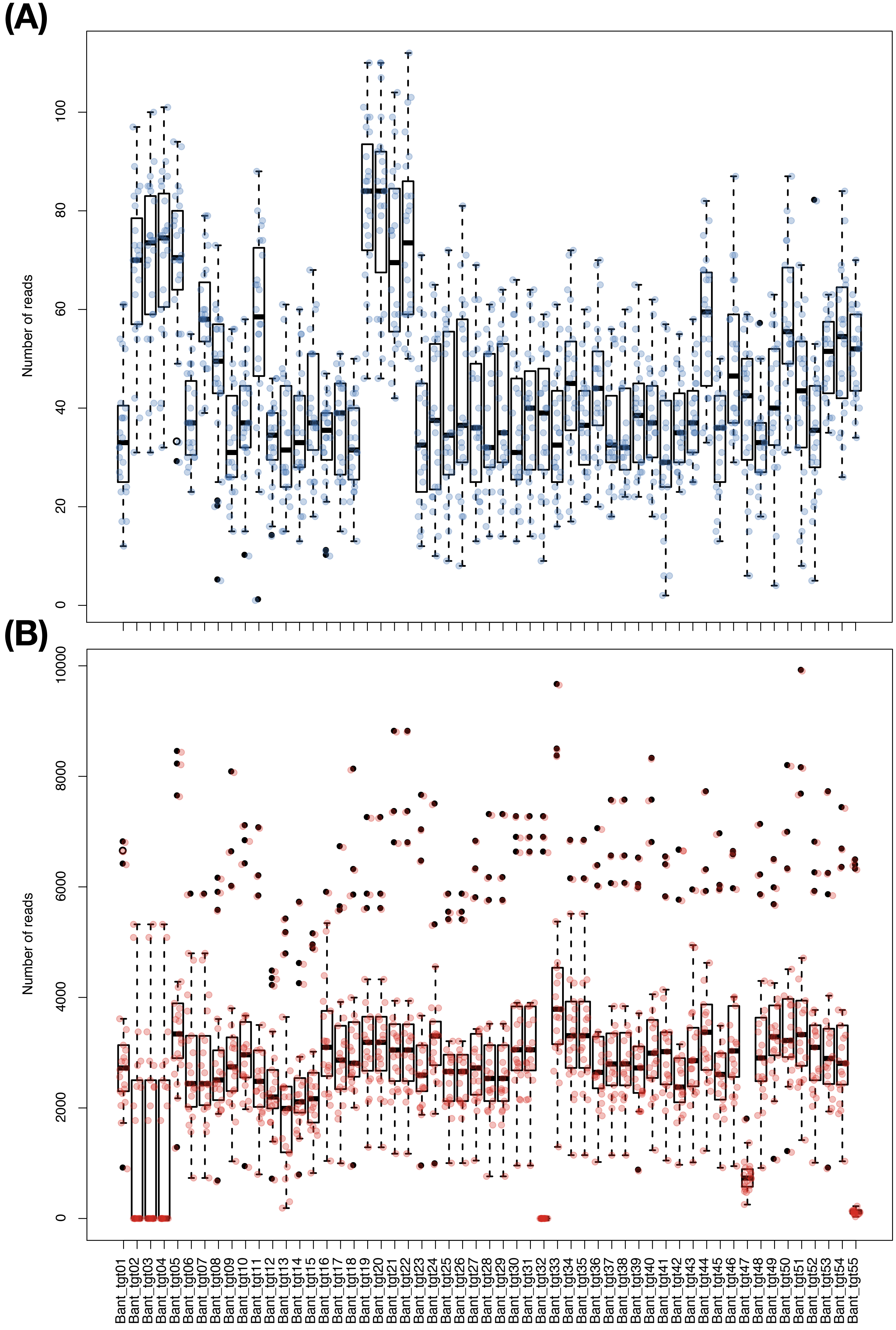

(A)
Number of reads
0
20
40
60
80
100
(B)
Number of reads
0
2000
4000
6000
8000
10000
Bant_tgt01
Bant_tgt02
Bant_tgt03
Bant_tgt04
Bant_tgt05
Bant_tgt06
Bant_tgt07
Bant_tgt08
Bant_tgt09
Bant_tgt10
Bant_tgt11
Bant_tgt12
Bant_tgt13
Bant_tgt14
Bant_tgt15
Bant_tgt16
Bant_tgt17
Bant_tgt18
Bant_tgt19
Bant_tgt20
Bant_tgt21
Bant_tgt22
Bant_tgt23
Bant_tgt24
Bant_tgt25
Bant_tgt26
Bant_tgt27
Bant_tgt28
Bant_tgt29
Bant_tgt30
Bant_tgt31
Bant_tgt32
Bant_tgt33
Bant_tgt34
Bant_tgt35
Bant_tgt36
Bant_tgt37
Bant_tgt38
Bant_tgt39
Bant_tgt40
Bant_tgt41
Bant_tgt42
Bant_tgt43
Bant_tgt44
Bant_tgt45
Bant_tgt46
Bant_tgt47
Bant_tgt48
Bant_tgt49
Bant_tgt50
Bant_tgt51
Bant_tgt52
Bant_tgt53
Bant_tgt54
Bant_tgt55

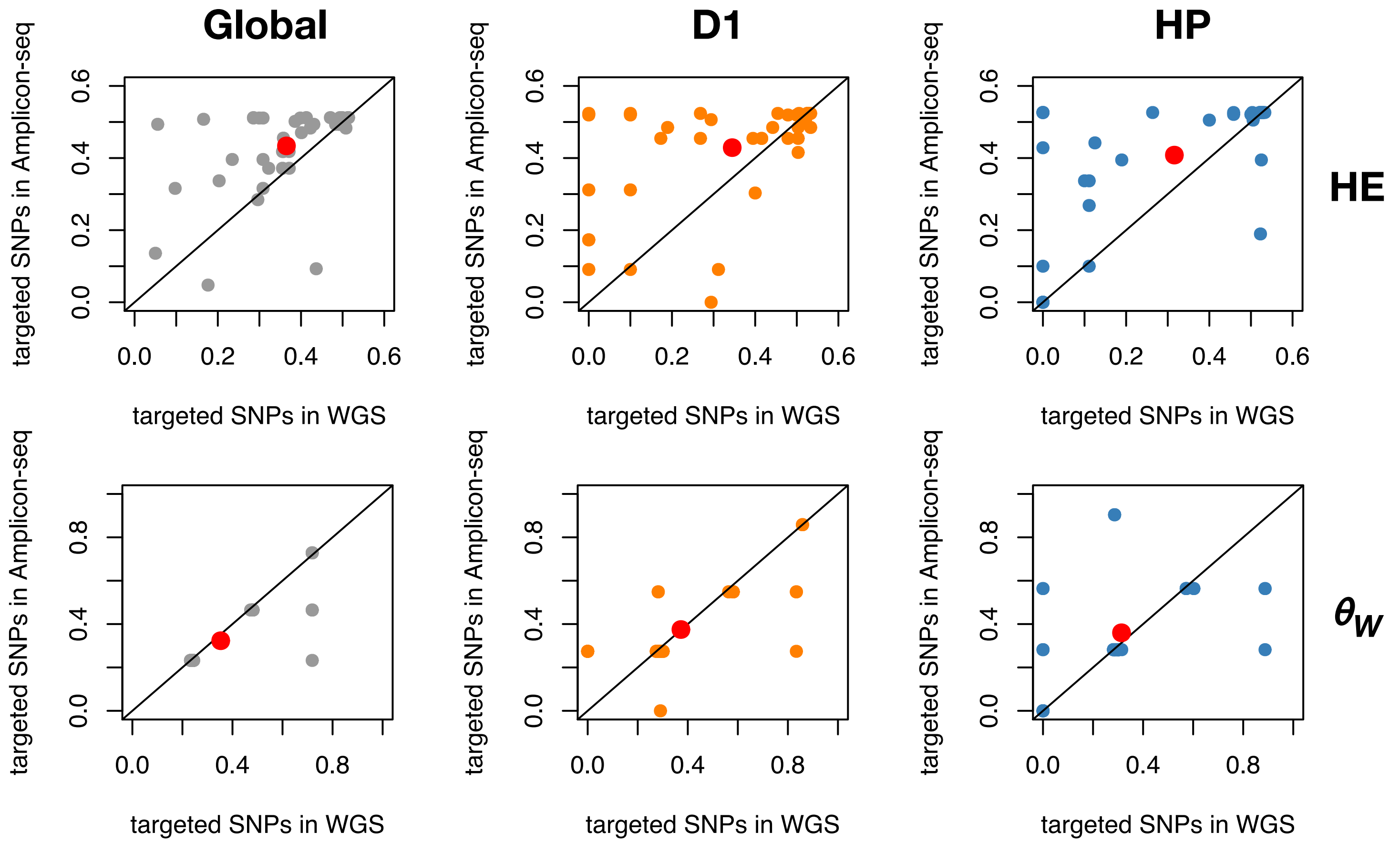
Global
D1
HP
HE
$\theta_W$
targeted SNPs in Amplicon-seq
targeted SNPs in WGS

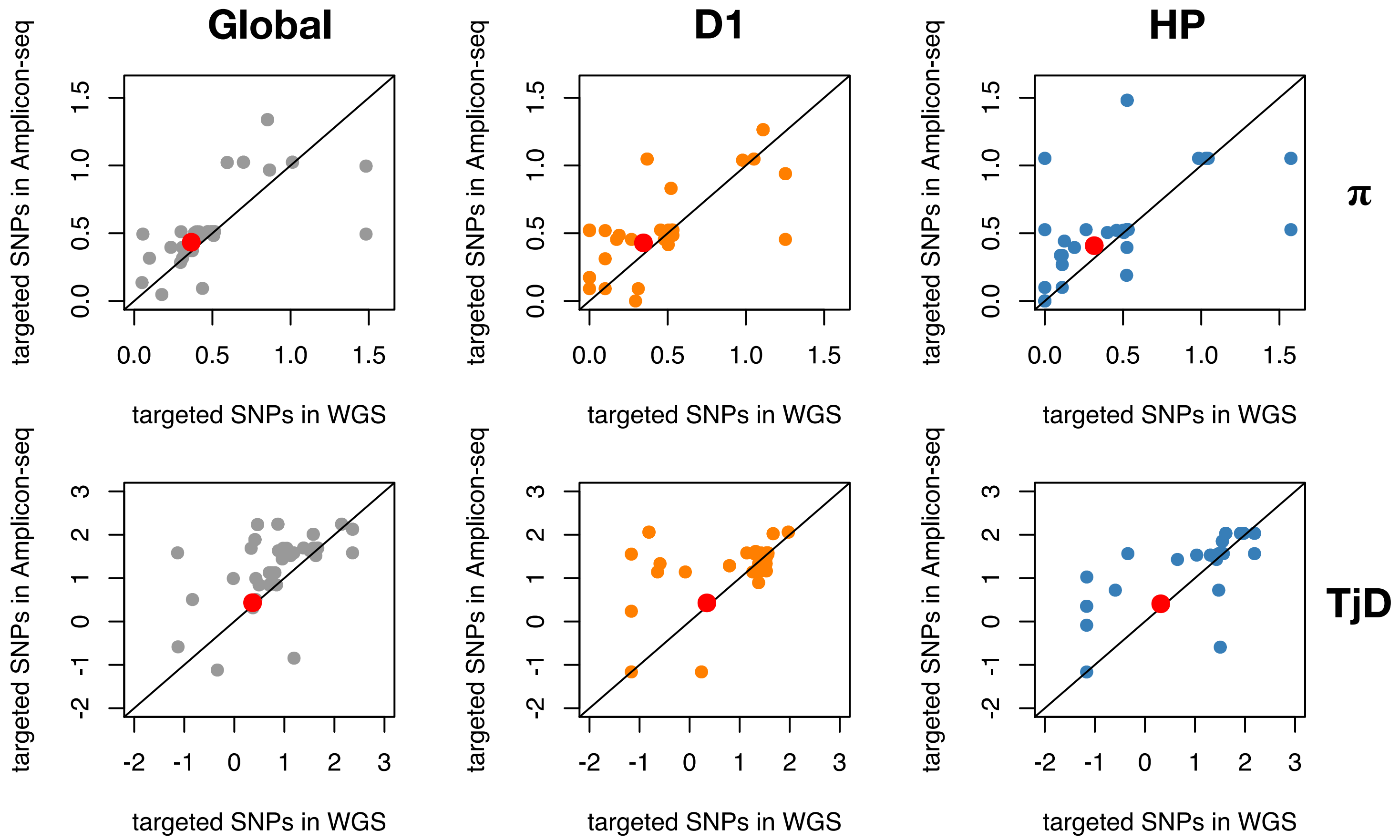
Global
D1
HP
π
TjD
targeted SNPs in Amplicon-seq
targeted SNPs in WGS
0.0
0.5
1.0
1.5
-2
-1
0
1
2
3

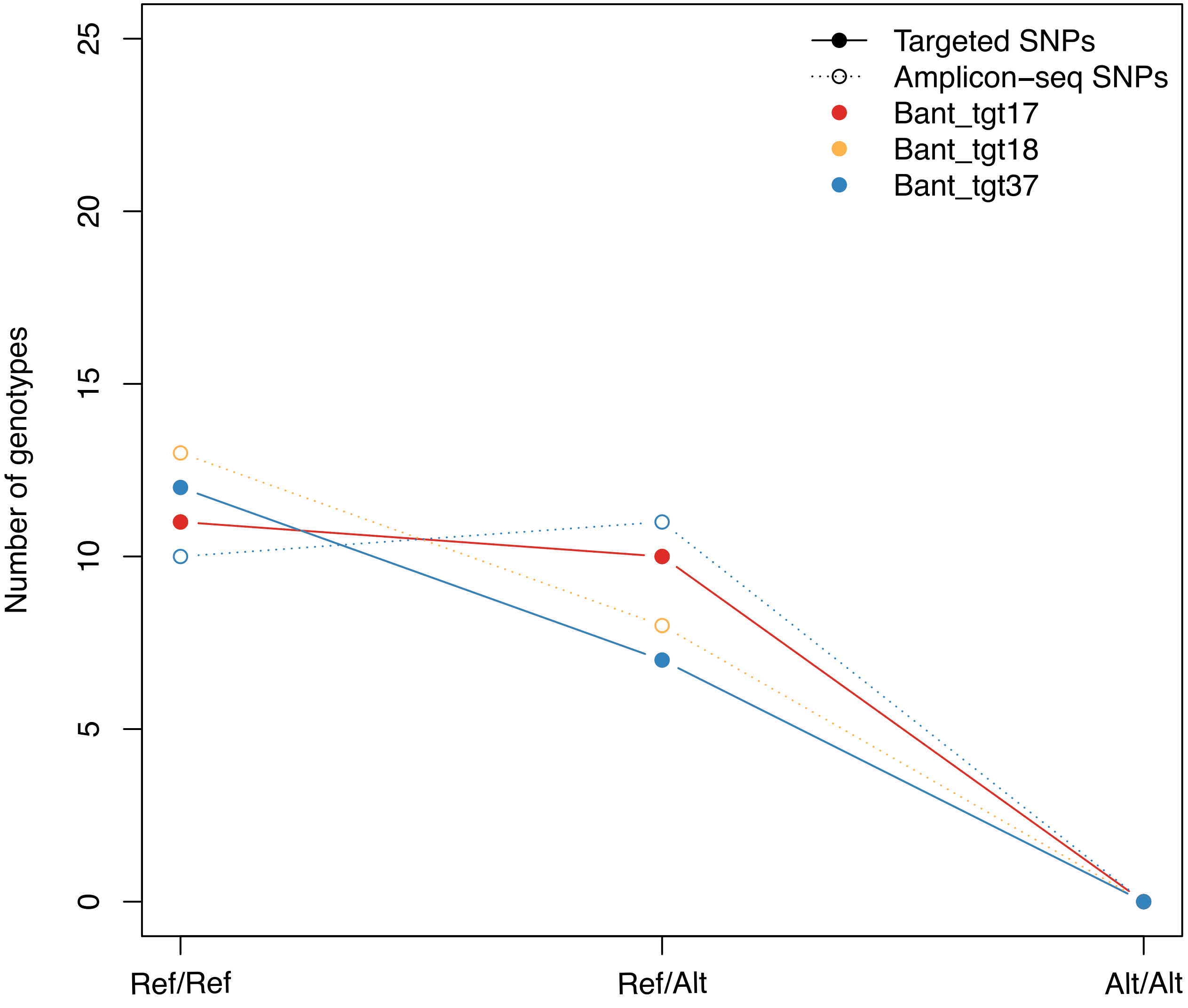
Targeted SNPs
Amplicon−seq SNPs
Bant_tgt17
Bant_tgt18
Bant_tgt37
Number of genotypes
0
5
10
15
20
25
Ref/Ref
Ref/Alt
Alt/Alt
